# Membrane Filtration Removes Lipids and Compromises the Acoustic Stability of Nanobubbles

**DOI:** 10.64898/2026.10.04.756539

**Authors:** Mahshid Yaali, Elizabeth Berndl, Aaqib Khan, Eric Strohm, Theresa Kosmides, Agata Exner, Michael Kolios

## Abstract

Nanobubbles (NBs) have been widely investigated as ultrasound contrast and therapeutic agents, yet whether submicron gas bubbles can directly generate measurable ultrasound signals remains controversial. Because acoustic scattering scales with bubble size, several studies have recently attributed nanobubble echogenicity primarily to larger bubbles. A recent Science Advances publication, attributed the loss of ultrasound signal after filtration to the removal of residual larger bubbles [1]. This interpretation assumes that filtration acts solely as a size-selection process. Here, we show that filtration also alters the shell properties and stability of lipid-coated NBs, independent of size. Filtered NBs (F-NBs) show significant lipid loss and accumulation on the filters themselves, resulting in decreased bubble stability and greater susceptibility to destruction under acoustic excitation. To decouple size from processing effects, we generated NB populations size-matched to filtered samples using increased centrifugation (IC-NBs), a comparatively gentler separation method. Despite comparable size distributions, IC-NBs retained sustained ultrasound scattering, whereas filtered NBs showed significantly reduced acoustic persistence. Filtration through 5 µm pores, ∼5-fold larger than the largest bubbles, preserved acoustic persistence but still reduced bubble concentration by ∼30%, suggesting that membrane contact may contribute to bubble loss beyond size exclusion. Complementary lipid and gas-content assays provided further evidence of filtration-induced changes. The 0.45 µm filtration removed substantially more total lipid than could be attributed to free-lipid removal alone, while the lipid associated with the larger bubbles removed during filtration accounted for only ∼5% of the additional lipid loss. Encapsulated gas content remained comparable to that of size-matched centrifuged controls, suggesting that the reduced acoustic stability of filtered NBs is more closely associated with changes in lipid content than with loss of the gas core. These results indicate that membrane filtration is not a purely size-selective process: it removes substantially more lipid than size exclusion predicts and yields bubbles that are less stable under ultrasound, most likely through changes in shell lipid content or in the surrounding lipid reservoir, underscoring the importance of processing-induced effects in interpreting nanobubble echogenicity.

## Introduction

Lipid-coated gas nanobubbles (NBs), typically defined as submicron gas-filled colloidal structures (also referred to as ultrafine bubbles [2]) have emerged as promising ultrasound contrast agents with applications in molecular imaging, drug delivery, and image-guided therapy [3], [4]. Their reduced size relative to conventional microbubbles enables prolonged circulation and potential access to extravascular compartments, motivating their development as next-generation theranostic agents [5], [6], [7]. Advances in formulation strategies have further enabled control over shell composition and interfacial properties to improve colloidal stability and functional performance under physiological conditions [6].

Despite substantial progress in formulation and characterization, whether nanobubble populations contribute directly to ultrasound contrast remains unresolved [8]. Classical bubble dynamics predicts a strong decrease in acoustic scattering with decreasing bubble radius, implying that submicron bubbles should generate weak echogenicity under diagnostic ultrasound conditions [9], [10]. In contrast, several experimental studies have reported measurable ultrasound signals from nanobubble formulations, leading to ongoing debate regarding the physical origin of the observed echogenicity [3], [4], [11], [12], [13]. A key unresolved question is therefore whether the measured signal arises from intrinsically active nanobubbles or larger bubbles (either the nanobubble size-distribution tail (400–800 nm) or residual populations of microbubbles ≥1 µm diameter).

A central difficulty in resolving this discrepancy is the inherently polydisperse nature of mechanically agitated bubble preparations. Standard fabrication methods produce broad size distributions spanning both nano- and microscale populations, even after conventional post-processing steps [9], [14]. Because acoustic scattering scales strongly with bubble radius, it is generally assumed that even trace amounts of micron-sized bubbles can disproportionately dominate the measured ultrasound response. Consequently, size-fractionation techniques such as differential centrifugation and membrane filtration are routinely employed to enrich submicron populations and reduce contributions from larger bubbles, although complete elimination of the upper size tail (400-800nm) is difficult to achieve [13], [15]. These procedures are typically assumed as passive separation methods that preserve the intrinsic physical properties of the remaining nanobubbles [1].

Recent experimental evidence indicates that removal of larger bubbles through filtration can eliminate measurable ultrasound signal, suggesting a strong contribution from size-dependent populations [1]. At the same time, this raises a critical question: whether the observed reduction in acoustic activity reflects only changes in size distribution or also alterations in bubble integrity induced during processing.

This question is particularly relevant for lipid-coated bubbles, whose acoustic response is governed not only by geometric size but also by shell elasticity, shell friction, permeability, and interfacial molecular organization [16], [17], [18], [19], [20], [21], [22]. Variations in lipid composition have been shown to alter nonlinear oscillation behavior, resonance frequency, and stability under acoustic excitation [21], [22], [23]. These effects arise because lipid monolayers are mechanically sensitive interfaces that can undergo stress-induced restructuring, phase transitions, or loss of structural integrity during processing steps such as filtration or high-speed centrifugation, potentially modifying bubble behavior independently of size distribution.

Here, we hypothesize that filtration and differential centrifugation differ in their effects on lipid-coated nanobubbles and influence physicochemical and acoustic properties in addition to size distribution. Specifically, we propose that reductions in acoustic activity after filtration arise not only from removing larger bubbles, but also from filtration-induced changes in shell integrity and mechanical stability. To test this hypothesis, we compare mechanically filtered nanobubbles with size-matched populations generated via increased centrifugation, a comparatively less disruptive separation method, using three independent replicates (bubble samples were independently prepared beginning at mechanical amalgamation from a stock lipid dispersion). We further investigate the effect of filtration under conditions where pore sizes exceed nominal bubble diameters, allowing separation of mechanical stress effects from size exclusion. By decoupling processing-induced effects from size distribution, this study directly tests a key assumption underlying current interpretations of nanobubble echogenicity and provides new insight into the physical determinants governing the acoustic response of lipid-coated bubbles.

## Results

### Filtration and increased centrifugation generate nanobubble populations with matched size distributions but divergent concentrations

To determine whether membrane filtration acts as a purely size-selective process, we generated five nanobubble (NB) populations from a common parent suspension: NBs isolated by standard differential centrifugation at 50 RCF (standard operating protocol (SOP) NB), SOP NBs passed through 0.45 μm or 5 μm polyvinylidene fluoride (PVDF) syringe filters (F-NB-0.45 and F-NB-5, respectively), and NBs isolated by increased-speed differential centrifugation at 100 or 150 RCF (IC-NB-100 and IC-NB-150), a mechanical method of narrowing the size distribution without direct exposure of bubbles to a membrane. The 5 μm filter pore size was chosen to be roughly five times larger than the largest bubbles present in the standard NB suspension, allowing us to test whether filtration produces effects independent of size exclusion.

All five bubble populations were characterized using three independent sample preparations by resonant mass measurement (RMM), dynamic light scattering (DLS), and nanoparticle tracking analysis (NTA); results from the three modalities were in close agreement (Figure 1). RMM was prioritized for interpretation because, unlike DLS and NTA, it distinguishes positively buoyant, gas-filled particles (NBs) from negatively buoyant lipid vesicles and micelles, although its nano-sensor is limited to particles from 0.1 to 2 μm [24]. DLS and NTA extended size characterization from 0.01 μm up to 15 μm, with no particles above ∼1 µm detected in any population. For clarity, Figure 1 focuses on the SOP NBs alongside F-NB-0.45 and IC-NB-150, the two size-matched populations most directly compared throughout this study, with the x-axis restricted to 1 μm to better resolve differences in curve shape between samples; the full five-population, three modality comparison, together with the extended size range confirming the absence of any larger microbubble population, is provided in Fig. S1.

**Figure 1.**
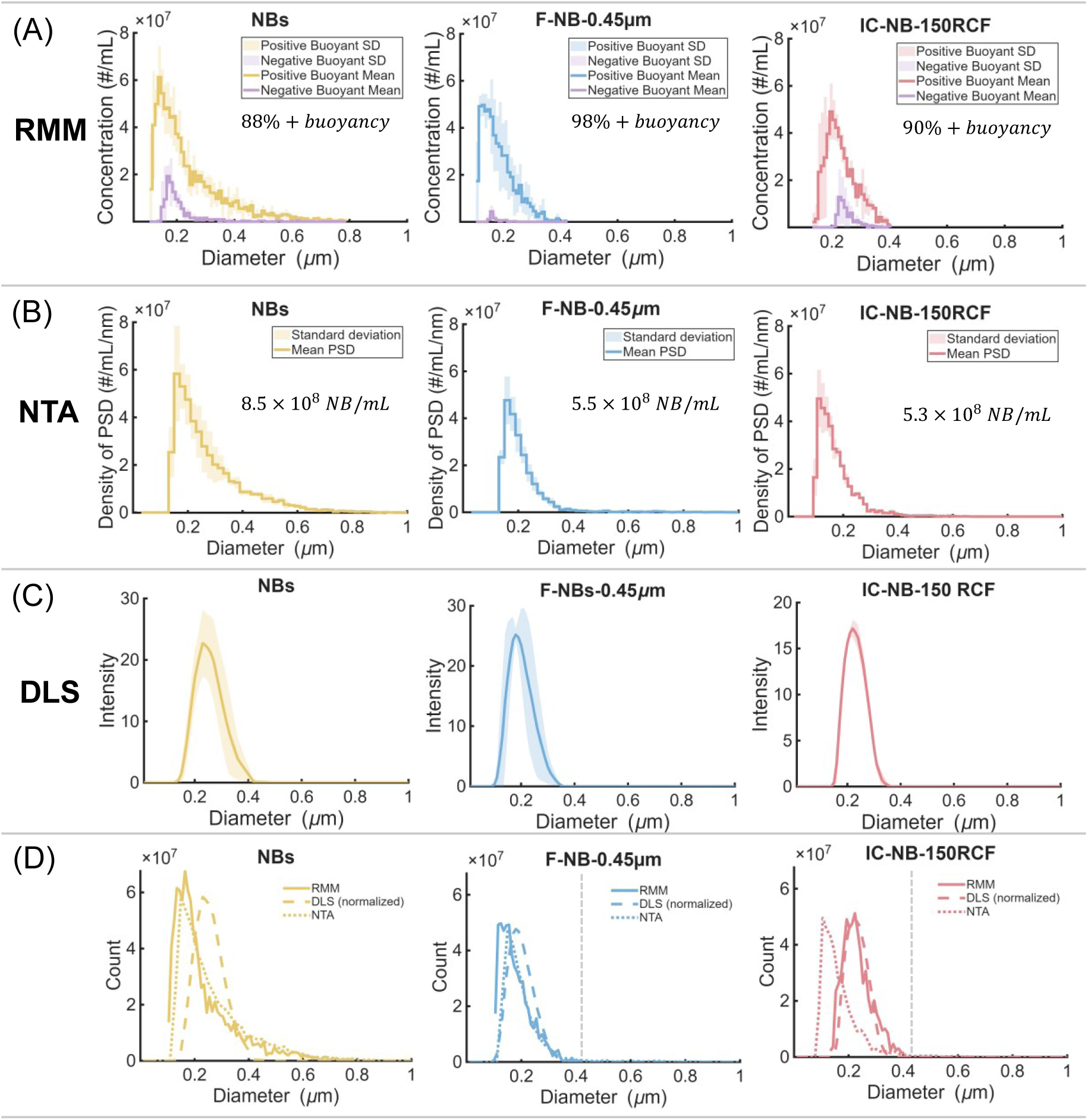
Size distribution and concentration characterization of SOP nanobubbles and two size-matched populations isolated by filtration or increased centrifugation. **(A)** Size distributions measured by resonant mass measurement (RMM) for SOP NBs (yellow), F-NB-0.45 (blue), and IC-NB-150 (red). Purple traces in all panels denote negatively buoyant (free lipid/liposomal) particles, shown for each sample. Each curve represents the mean of n = 3 independent trials; shaded regions indicate SD. **(B)** Nanoparticle tracking analysis (NTA): SOP NBs show the highest bubble concentration, with sizes spanning up to approximately 800 nm within this 1 µm window. F-NB-0.45 and IC-NB-150 show closely matched, narrower size distributions truncated near 400 nm, accompanied by IC-NB-150 retained significantly greater bubble concentrations 35% and 37% lower than SOP NBs, respectively, despite arising from two mechanistically distinct isolation methods. and **(C)** dynamic light scattering (DLS) size distributions for the same three samples (n = 3 each), confirming the size characteristics observed by RMM. **(D)** Mean size distributions from all three modalities overlaid for each sample (n = 3 per modality). Each modality has its own detection limits and sensitivity, which is reflected in differences in the mean curves; however, RMM and NTA in particular show close agreement in the upper size tail of each population, indicating consistent estimates of the largest particles present and, correspondingly, of the size-isolation efficiency of each method. The x-axis is restricted to 1 µm across all panels to better resolve differences in curve shape between samples. A gray dotted line marks the 0.45 µm cut-off, allowing visual comparison of how effectively each isolation method excludes larger particles; the full five-population comparison (including F-NB-5 and IC-NB-100), extended to 15 µm to confirm the absence of any larger microbubble population, is provided in Fig. S1.

As expected, F-NB-5 and SOP NBs exhibited similar size distributions, with both showing an upper tail extending to approximately 800 nm (Fig. S1). In contrast, F-NB-0.45 and IC-NB-150 showed comparable distributions with the upper tail truncated near 400 nm (Fig. 1A), whereas IC-NB-100 exhibited a slightly broader distribution extending to approximately 500 nm (Fig. S1). This design yields two independent routes to a comparable size-restricted population, mechanical filtration (F-NB-0.45) versus centrifugal sedimentation (IC-NB-150), allowing size and processing method to be decoupled to the extent that the measured distributions are comparable (Figure 1). Because acoustic scattering is strongly weighted toward the largest bubbles, small differences in the upper tail (∼350–450 nm) are not resolved by these measurements and could contribute to differences in signal.

Concentration, however, did not track simply with pore size. Despite a pore diameter roughly five-fold larger than the largest bubbles present, F-NB-5 showed a 30% reduction in bubble concentration relative to SOP NBs (Fig. S1). F-NB-0.45 showed a comparable 35% reduction (Figure 1A). Increased centrifugation bubbles had 18% lower bubble concentration than the SOP isolate for IC-NB-100 (Fig. S1), and IC-NB-150 showed a 37% lower concentration (Figure 1A), more similar in magnitude to the 30% and 35% reductions produced by filtration. The fact that a 5 µm filter, nominally too large to physically exclude any bubble in the sample, still removed nearly a third of the population is a first indication that filtration is not acting solely through size exclusion.

### Size alone does not explain the loss of ultrasound contrast in filtered nanobubbles

We next compared the acoustic response of three independent preparations across all five populations within agarose phantoms using concurrent B-mode and contrast-mode imaging at 21 MHz over 300 frames with a preclinical small-animal ultrasound system (FUJIFILM VisualSonics VEVO 2100). SOP NBs and IC-NBs were diluted to a matched concentration of 1 × 10⁹ NB mL^−1^ prior to imaging, whereas F-NBs were imaged at their post-filtration concentrations (Methods). At moderate acoustic pressures (0.81 MPa B-mode; 0.62 MPa contrast mode), SOP NBs and F-NB-5 produced comparable, stable signal amplitudes above PBS throughout the imaging window (Figure 2A, C). F-NB-0.45, in contrast, produced a signal slightly above PBS for approximately 120 s in B-mode and showed no measurable increase over PBS in contrast mode. When signal amplitude was time-averaged across all 300 frames for each batch and normalized to PBS (Figure 2B, D), SOP NBs and F-NB-5 were significantly higher than both PBS and F-NB-0.45 in B-mode and contrast mode (p < 0.0001 for each comparison; one-way ANOVA with Tukey HSD, n = 3 independent batches). F-NB-5 did not differ significantly from SOP NBs in either mode (p > 0.05), whereas F-NB-0.45 was not statistically different from PBS in either mode (p > 0.99).

**Figure 2.**
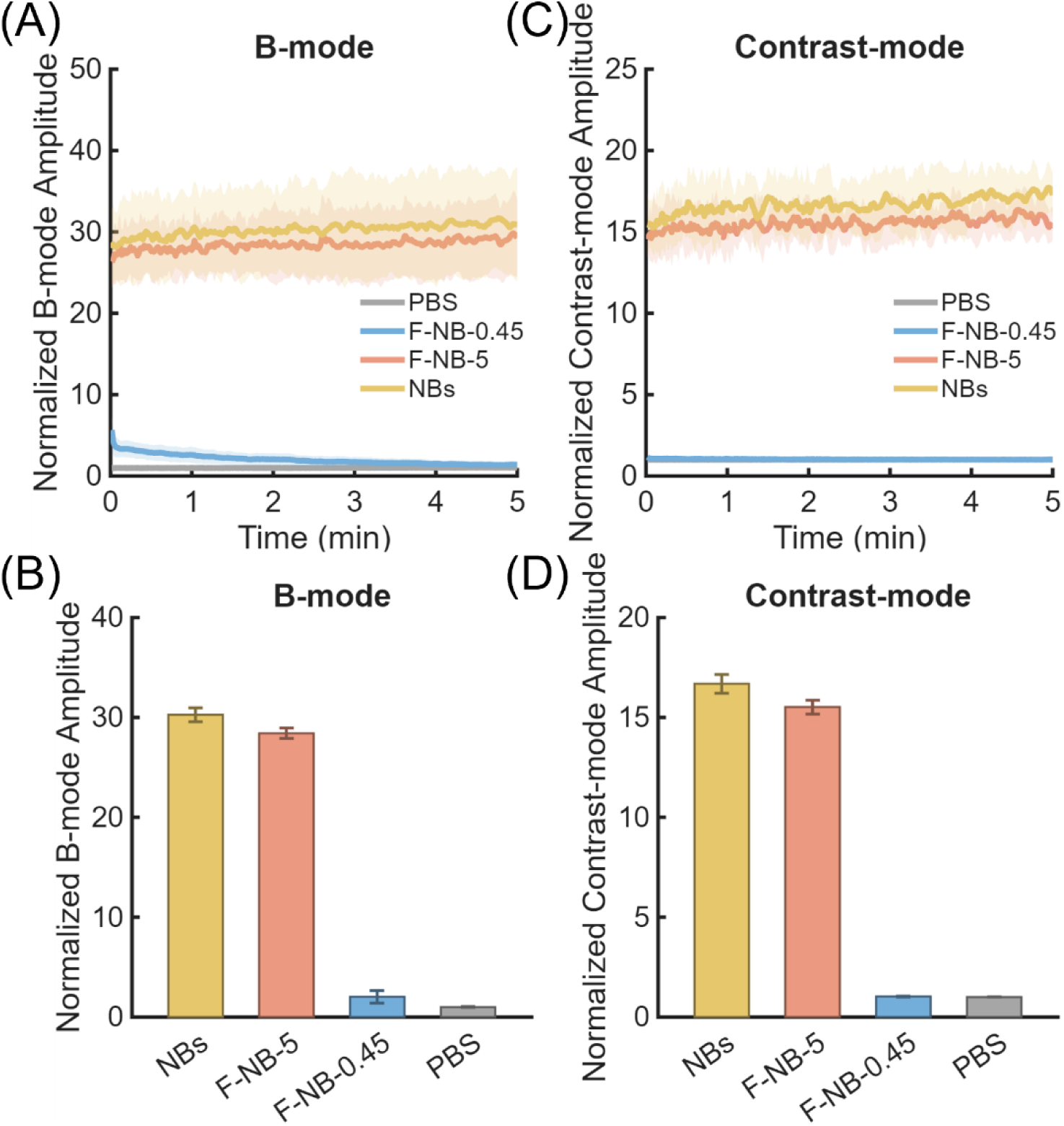
Ultrasound contrast response of SOP and filtered nanobubbles at moderate acoustic pressure. **(A)** B-mode (0.81 MPa) normalized signal amplitude over 300 s for SOP NBs, F-NB-5, F-NB-0.45, and a PBS control, acquired using a 21 MHz linear array transducer. **(B)** Mean B-mode signal amplitude (time-averaged over all 300 frames for each batch), normalized to PBS, for the same samples. **(C)** Contrast-mode (0.62 MPa) normalized signal amplitude over 300 s for the same samples. **(D)** Mean contrast-mode signal amplitude (time-averaged over all 300 frames for each batch), normalized to PBS. F-NB-0.45 showed a transient signal slightly above PBS for approximately 120 s in B-mode only (A) and no measurable increase over PBS in contrast mode (Contrast-mode (0.62 MPa) normalized signal amplitude over 300 s for the same samples. **(D)** Mean contrast-mode signal amplitude (time-averaged over all 300 frames for each batch), normalized to PBS. F-NB-0.45 shows signal above PBS in B-mode only (A) and no measurable increase over PBS in contrast mode (C). In the time-averaged data (B, D), SOP NBs and F-NB-5 were significantly higher than both PBS and F-NB-0.45 (p < 0.0001 for each comparison), whereas F-NB-5 was not significantly different from SOP NBs (p > 0.05). F-NB-0.45 was not significantly different from PBS in either mode (p > 0.99). Statistics: one-way ANOVA with Tukey HSD on batch-level values. Data represent n = 3 independent batches per group.

To distinguish the effects of size and bubble concentration, F-NB-0.45 was compared with IC-NB-150, which had a similar upper size limit of approximately 400 nm but was generated without filter-membrane contact (Figure 3). Although IC-NB-150 was imaged at approximately 1.5-fold higher bubble concentration (1 × 10⁹ versus ∼0.65 × 10⁹ NB mL^−1^), this difference in bubble number cannot account for the approximately eightfold higher signal observed for IC-NB-150 relative to F-NB-0.45. Moreover, F-NB-5 retained SOP-level acoustic signal despite a ∼30% reduction in bubble number. Together, these comparisons indicate that neither differences in bubble number nor the reduction in upper bubble size alone can explain the loss of acoustic activity following 0.45 µm filtration.

**Figure 3.**
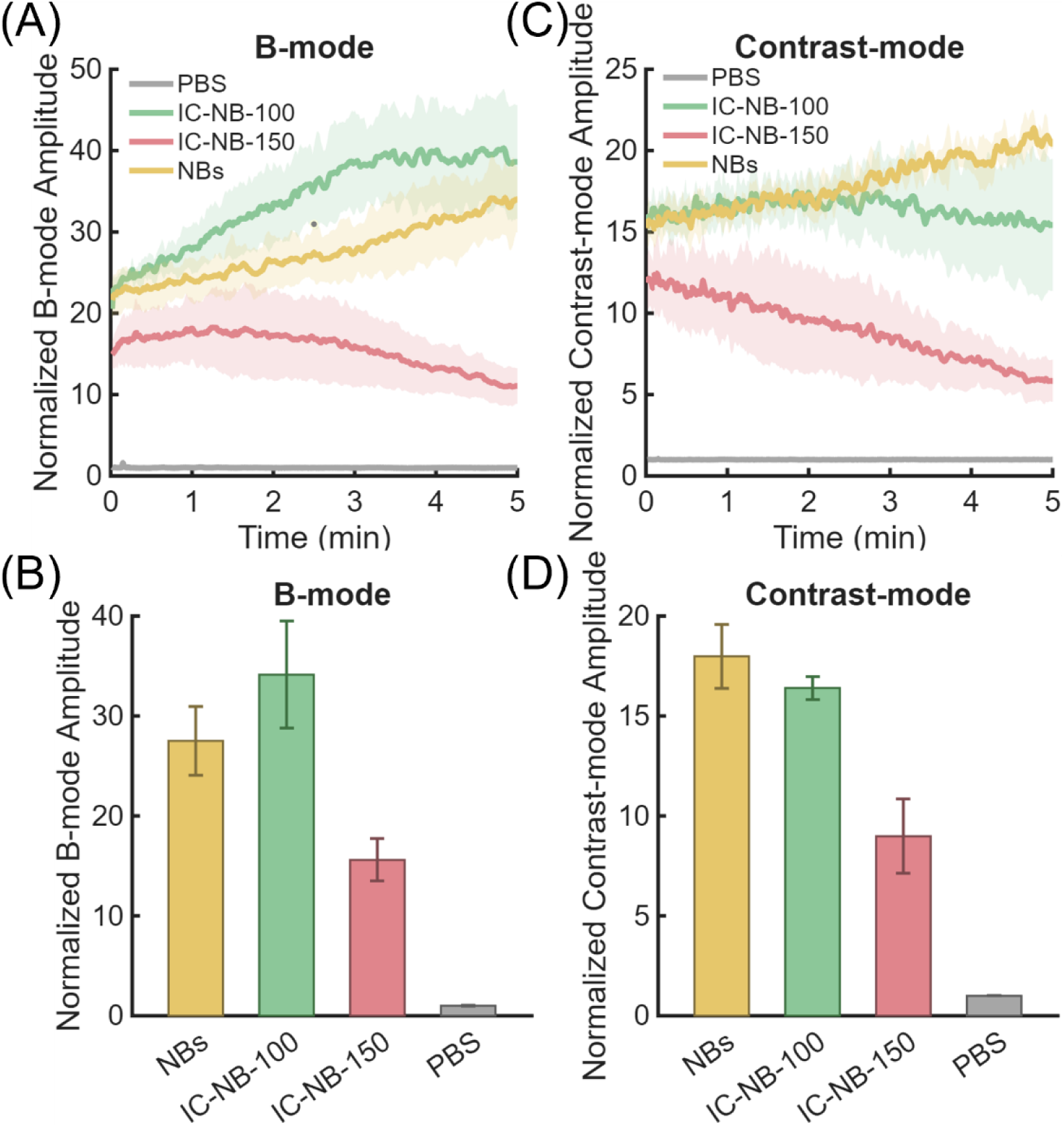
Ultrasound contrast response of size-matched centrifuged nanobubbles at moderate acoustic pressure. **(A)** B-mode (0.81 MPa) normalized signal amplitude over 300 s for SOP NBs (50 RCF), IC-NB-100, IC-NB-150, and PBS. **(B)** Mean B-mode signal amplitude, time-averaged over the full 300 s for each batch and normalized to PBS. **(C)** Contrast-mode (0.62 MPa) normalized signal amplitude over 300 s for the same samples. **(D)** Mean contrast-mode signal amplitude, time-averaged over the full 300 s for each batch and normalized to PBS. All three centrifuged populations maintained measurable, comparatively stable signal throughout the acquisition (A, C) and showed mean amplitudes significantly above PBS (B, D; IC-NB-100 and SOP NBs, p < 0.0001; IC-NB-150, p = 0.002 in B-mode and p = 0.0004 in contrast mode). IC-NB-100 was not significantly different from SOP NBs (B, D; p = 0.23 and p = 0.80). IC-NB-150 showed a mean amplitude approximately 45–55% lower than IC-NB-100 and SOP NBs (B, D; p ≤ 0.010 for each comparison) but remained approximately 9-to 16-fold above PBS, in contrast to the size-comparable F-NB-0.45 population, whose signal was indistinguishable from PBS (p > 0.99). Statistics: one-way ANOVA with Tukey HSD on batch-level values. Data represent n = 3 independent batches per group.

This loss of contrast could, in principle, simply reflect the smaller size distribution of F-NB-0.45. To test this, we compared the acoustic signal from F-NB-0.45 against IC-NB-150, a population matched in size but generated without filter membrane contact (Figure 3). Unlike F-NB-0.45, both IC-NB-100 and IC-NB-150 produced signal significantly above PBS in B-mode (p < 0.0001 and p = 0.002) and contrast mode (p < 0.0001 and p = 0.0004; one-way ANOVA with Tukey HSD, n = 3 independent batches). Across the imaging window, all three centrifuged populations (50 (SOP), 100, and 150 RCF) maintained measurable, comparatively stable signal (Figure 3A, C). IC-NB-100 was statistically indistinguishable from SOP NBs in both modes (p = 0.23 and p = 0.80). IC-NB-150 showed reduced signal, roughly 45–55% lower than IC-NB-100 and SOP (B-mode: p = 0.0002 vs IC-NB-100, p = 0.010 vs SOP; contrast mode: p = 0.0008 vs IC-NB-100, p = 0.0001 vs SOP), consistent with the expected decrease in scattering with smaller bubble size. Critically, this reduced signal remained about 9-to 16-fold above PBS, whereas F-NB-0.45 was indistinguishable from PBS (p > 0.99 in both modes). IC-NB-150 retained significantly greater signal than F-NB-0.45 (B-mode p = 0.004; contrast mode p = 0.0004) despite comparable size distributions, indicating that the loss of acoustic activity following filtration cannot be explained by particle-size reduction alone. Nor can it be explained by bubble loss alone: F-NB-5 lost a comparable fraction of bubbles (30%) yet retained SOP-level signal, and a ∼35% reduction in bubble number cannot account for the complete loss of signal in F-NB-0.45. The difference instead points to the isolation method itself. This comparison establishes the central question addressed in the remainder of this study: what physically distinguishes F-NB-0.45 from IC-NB-150, two populations with matched size distributions but different acoustic behavior?

### Size-matched centrifuged nanobubbles withstand high-pressure ultrasound; filtered nanobubbles do not

To test whether this divergence extends to bubble survival under higher ultrasound pressure amplitude, we imaged F-NBs and IC-NBs at higher pressure (1.33 MPa B-mode, 1.18 MPa contrast mode) and compared the first and last acquired frames (Figure 4). F-NB-0.45 signal collapsed within the first few frames of exposure. F-NB-5 retained comparatively stable signal over the same interval. IC-NB samples matched in size to F-NB-0.45 showed markedly different behavior: both IC-NB-100 and IC-NB-150 maintained stable signal across the same high-pressure exposure and imaging duration (Figure 5). Thus, a nanobubble population with an identical size distribution to F-NB-0.45 survives high-pressure ultrasound when generated by centrifugation but is destroyed within seconds when generated by filtration, reinforcing that pore size alone does not determine bubble fragility.

**Figure 4.**
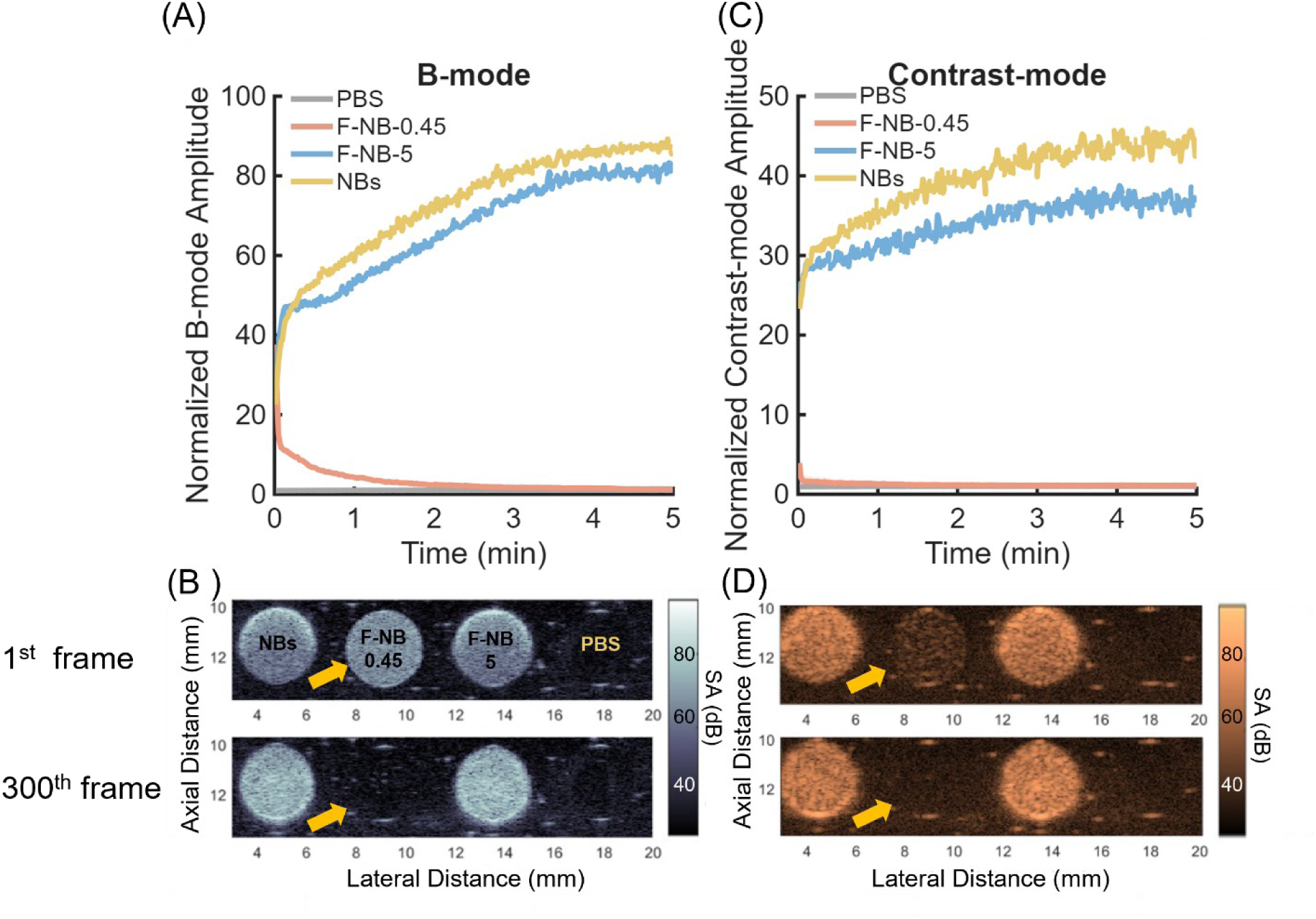
Filtered nanobubbles are destroyed by high-pressure ultrasound while SOP and large-pore-filtered nanobubbles remain stable. **(A)** B-mode (1.33 MPa) normalized signal amplitude over time for SOP NBs, F-NB-5, and F-NB-0.45. **(B)** Representative B-mode imaging frames for the same samples at the first (top) and 300th (bottom) frame of acquisition; arrows indicate the F-NB-0.45 well, highlighting the loss of signal by the final frame. **(C)** Contrast-mode (1.18 MPa) normalized signal amplitude over time for the same samples. **(D)** Representative contrast-mode imaging frames at the first (top) and 300th (bottom) frame; arrows indicate the F-NB-0.45 well. F-NB-0.45 signal collapses within the first few frames of exposure in both imaging modes, while SOP NBs and F-NB-5 retain comparatively stable signal across the full acquisition.

**Figure 5.**
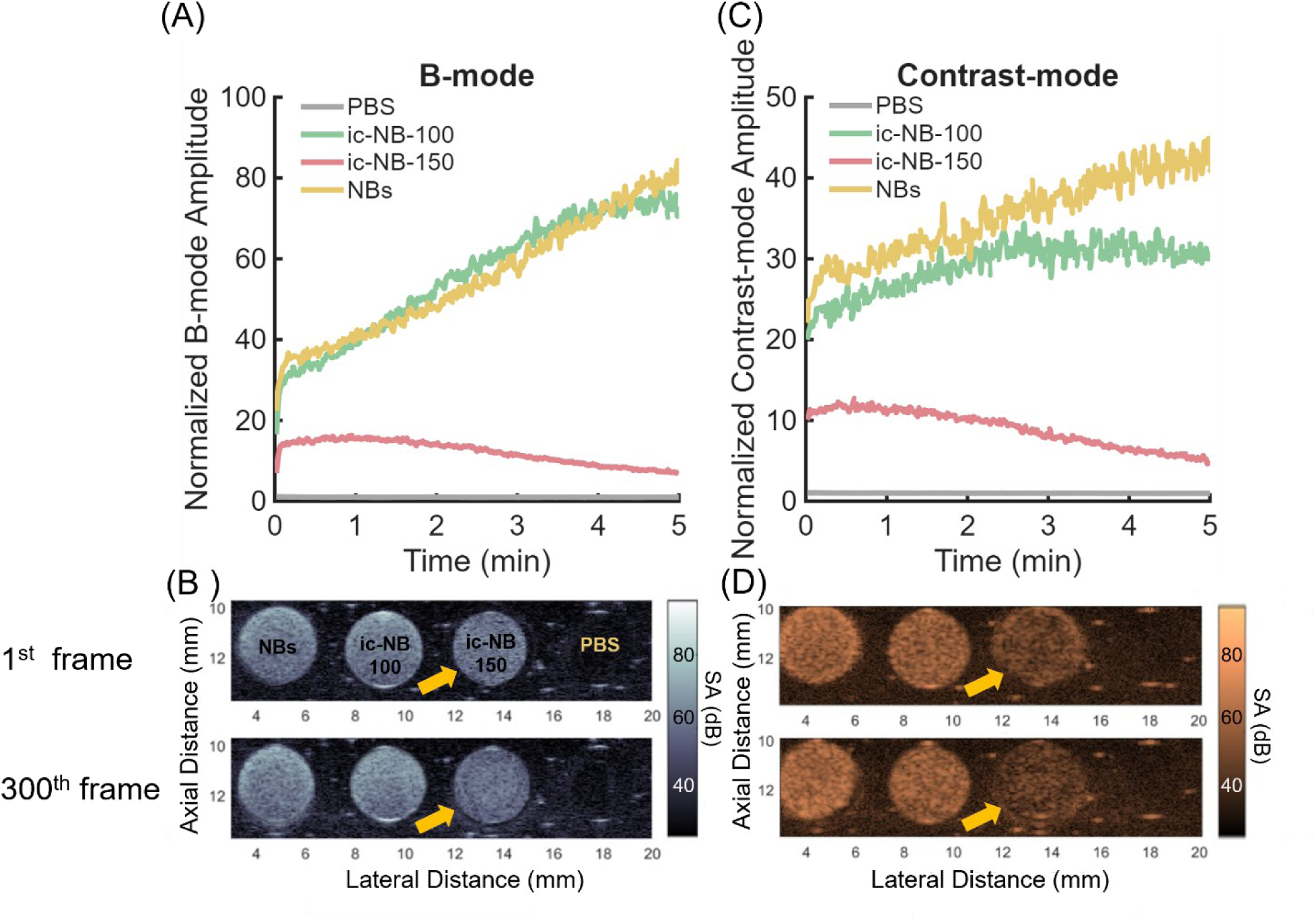
Size-matched increased-centrifugation nanobubbles withstand high-pressure ultrasound. **(A)** B-mode (1.33 MPa) normalized signal amplitude over time for SOP NBs, IC-NB-100, and IC-NB-150. **(B)** Representative B-mode imaging frames for the same samples at the first (top) and 300th (bottom) frame of acquisition; arrows indicate the IC-NB-150 well, highlighting that, unlike F-NB-0.45 (Figure 4), signal is retained by the final frame. **(C)** Contrast-mode (1.18 MPa) normalized signal amplitude over time for the same samples. **(D)** Representative contrast-mode imaging frames at the first (top) and 300th (bottom) frame; arrows indicate the IC-NB-150 well. Both IC-NB-100 and IC-NB-150, matched in size distribution to F-NB-0.45, maintain stable signal across the same high-pressure exposure and imaging duration under which F-NB-0.45 is destroyed.

### Filtration selectively depletes lipid from the nanobubble shell rather than removing only free lipid

Using the Stewart phospholipid assay, total recoverable lipid dropped from approximately 0.0174 mg to 0.0029 mg after 0.45 µm filtration, corresponding to an 83.3% reduction, compared with a more modest reduction to approximately 0.0095 mg (45.4%) after 5 µm filtration (Figure 6). In comparison, the total lipid content of the IC-NB-100 and IC-NB-150 samples was reduced to 0.015 mg (13.8%) and 0.0163 mg (6.3%), respectively, relative to the SOP NBs. The substantially greater lipid loss observed following membrane filtration, particularly with the 0.45 µm filter, is consistent with filtration-associated removal of lipid from the NB suspensions. However, this assay quantifies total phospholipid and cannot distinguish bubble-shell lipid from free lipid structures (micelles, liposomes, small vesicles).

**Figure 6.**
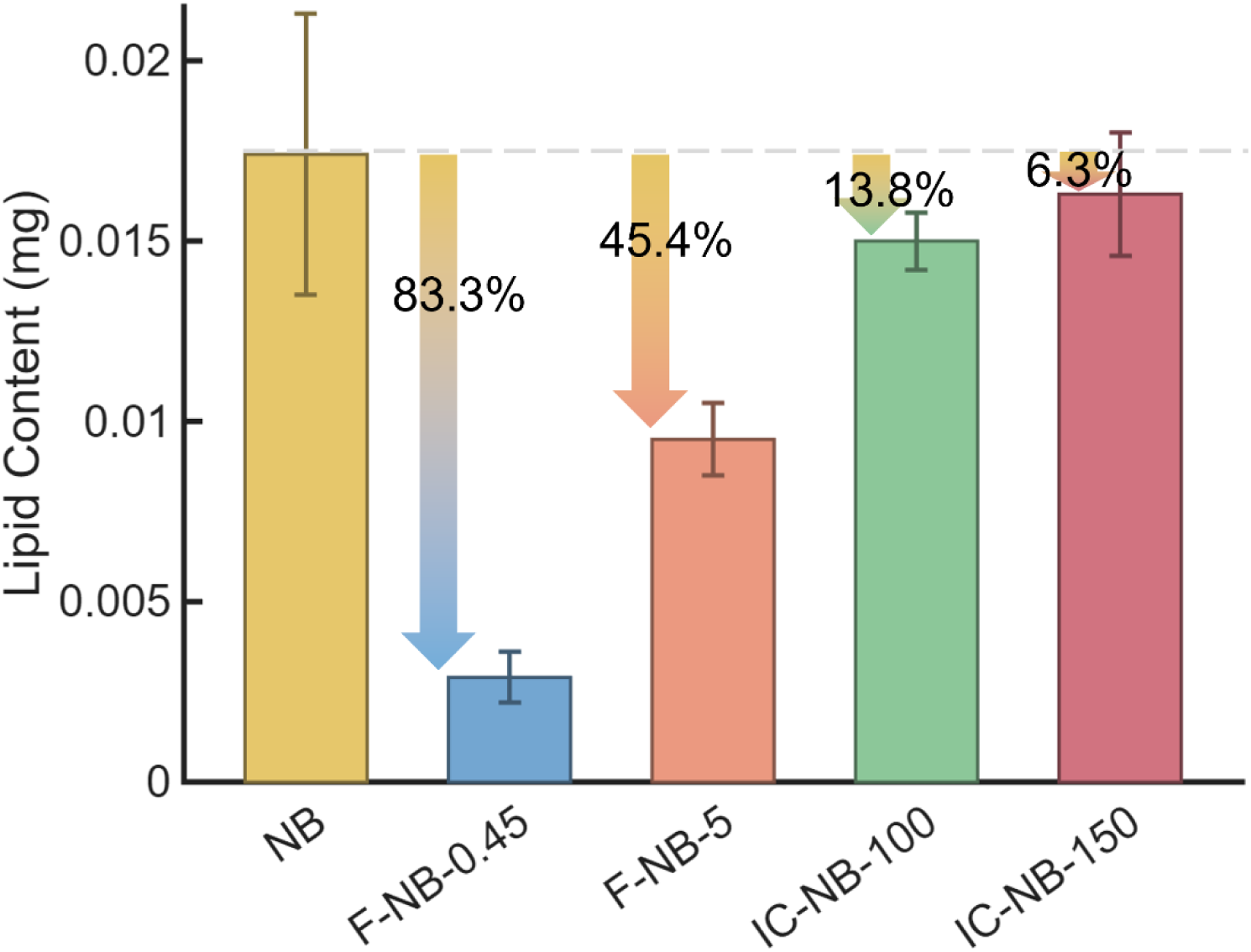
Membrane filtration results in greater loss of recoverable phospholipid than increased centrifugation. Total phospholipid content of SOP NBs filtered NB suspensions (F-NB-0.45 and F-NB-5), and centrifuged NB suspensions (IC-NB-100 and IC-NB-150), quantified using the Stewart ferrothiocyanate assay. Filtration through 0.45 µm and 5 µm PVDF membranes reduced total lipid content by 83.3% (from approximately 0.0174 mg to 0.0029 mg) and 45.4% (to approximately 0.0095 mg), respectively. In comparison, IC-NB-100 and IC-NB-150 showed reductions of 13.8% (to 0.015 mg) and 6.3% (to 0.0163 mg), respectively, relative to SOP NBs. Data are presented as mean ± SD (n = 3 independent batches per group).

To isolate the free-lipid contribution, we used RMM to separately quantify the negatively buoyant particle fraction, lipid vesicles and micelles that did not contribute to bubble formation, before and after filtration (Figure 7). After 5 µm filtration, the total lipid content decreases by 45.4%, while the free lipid fraction measured by RMM decreases by 33%, indicating that the majority of the lipid loss consists of free lipids. However, after 0.45um filtration, the total lipid content decreases by 83.3%, but the free lipid content decreases by only 42%. Free-lipid removal, as indexed by RMM-detectable vesicles, did not scale with the total lipid loss, suggesting that filtration also removes lipid associated with bubbles, either from removal of larger NBs or from the shells of survivors; resolving these contributions requires shell-specific measurements. Bubbles that survive filtration persist as gas-core particles and continue to be detected by particle-sizing methods (Figure 1) but are less stable under ultrasound exposure than size-matched centrifuged bubbles (Figures 2–5). The greater total lipid loss after 0.45 µm filtration is consistent with a difference in lipid content or shell integrity between F-NB-0.45 and IC-NB-150, although the present assays do not measure the shell directly.

**Figure 7.**
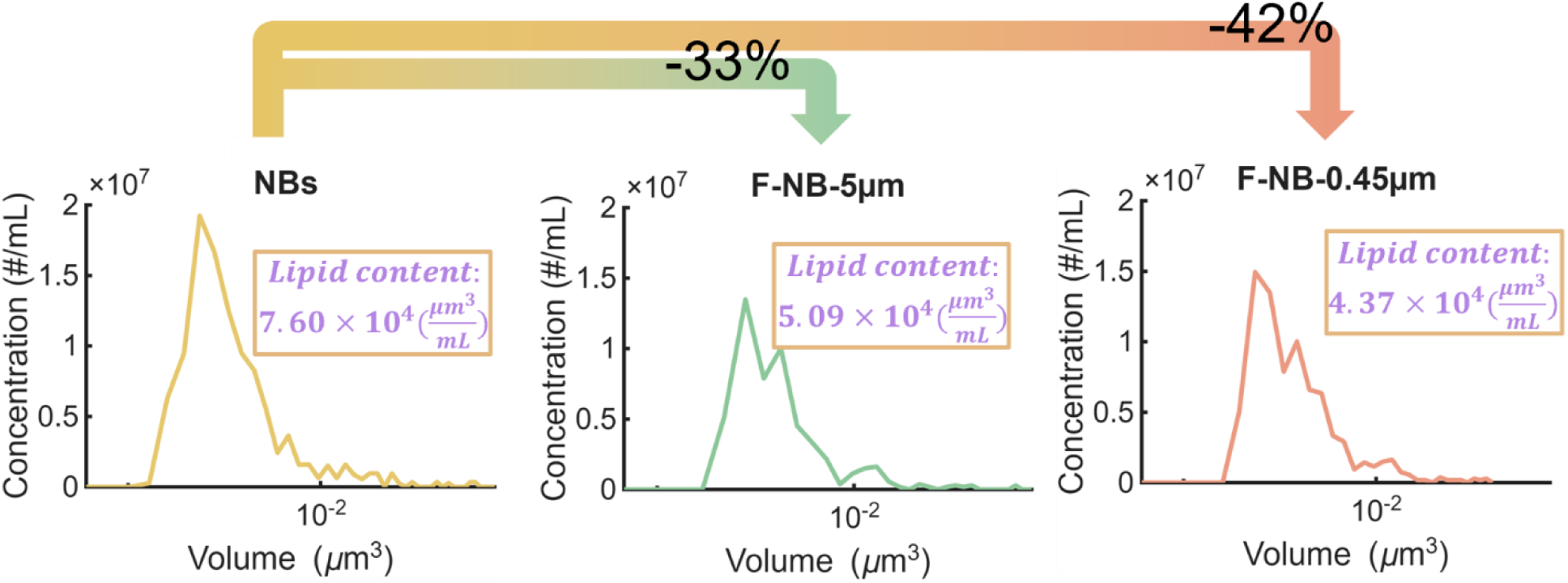
Filtration removes shell-associated lipid in addition to free lipid. Concentration of negatively buoyant particles (free lipid aggregates and liposomal material not associated with gas-filled nanobubbles) as a function of particle volume (logarithmic scale), measured by resonant mass measurement (RMM), before and after filtration through 0.45 µm and 5 µm membranes. Boxed values report the area under each curve, corresponding to the total volume of free lipid particles per mL; assuming constant lipid density across samples, fractional volume loss is equivalent to fractional mass loss. By this measure, free lipid content decreased by 42% and 33% for F-NB-0.45 and F-NB-5, respectively. This is substantially less than the corresponding total lipid mass loss measured by the Stewart assay (Figure 6) for the F-NB-0.45, indicating that a large fraction of the lipid removed by filtration originates from intact nanobubble shells rather than unincorporated free lipid. Data are shown as mean ± SD, n = 3 independent batches per group.

To test whether removal of larger bubbles could account for the additional lipid lost in F-NB-0.45, we estimated the shell lipid mass carried by the bubbles present in F-NB-5 but absent from F-NB-0.45 (Methods). Summed over the RMM difference distribution, this population contained 0.0003 mg of lipid on average, only ∼5% of the 0.0066 mg difference in total lipid between F-NB-5 and F-NB-0.45. Removal of the larger bubbles therefore explains little of the lipid deficit; the remaining ∼0.0063 mg must derive from lipid that is not part of the removed bubbles. The removed bubbles would need to carry ∼22-fold more lipid per bubble than a monolayer of their size to account for the difference. Filtration therefore removes far more lipid than the excluded bubbles carried, depleting the lipid available to the bubble shells. Together with the reduced acoustic stability of F-NB-0.45, this is consistent with filtration compromising shell integrity.

Optical microscopy of the filter membranes after use showed visible deposition of material on the membrane surface compared with unused membranes (Figure 8), which is consistent with, though not specific to, retention of lipid-containing material during filtration.

**Figure 8.**
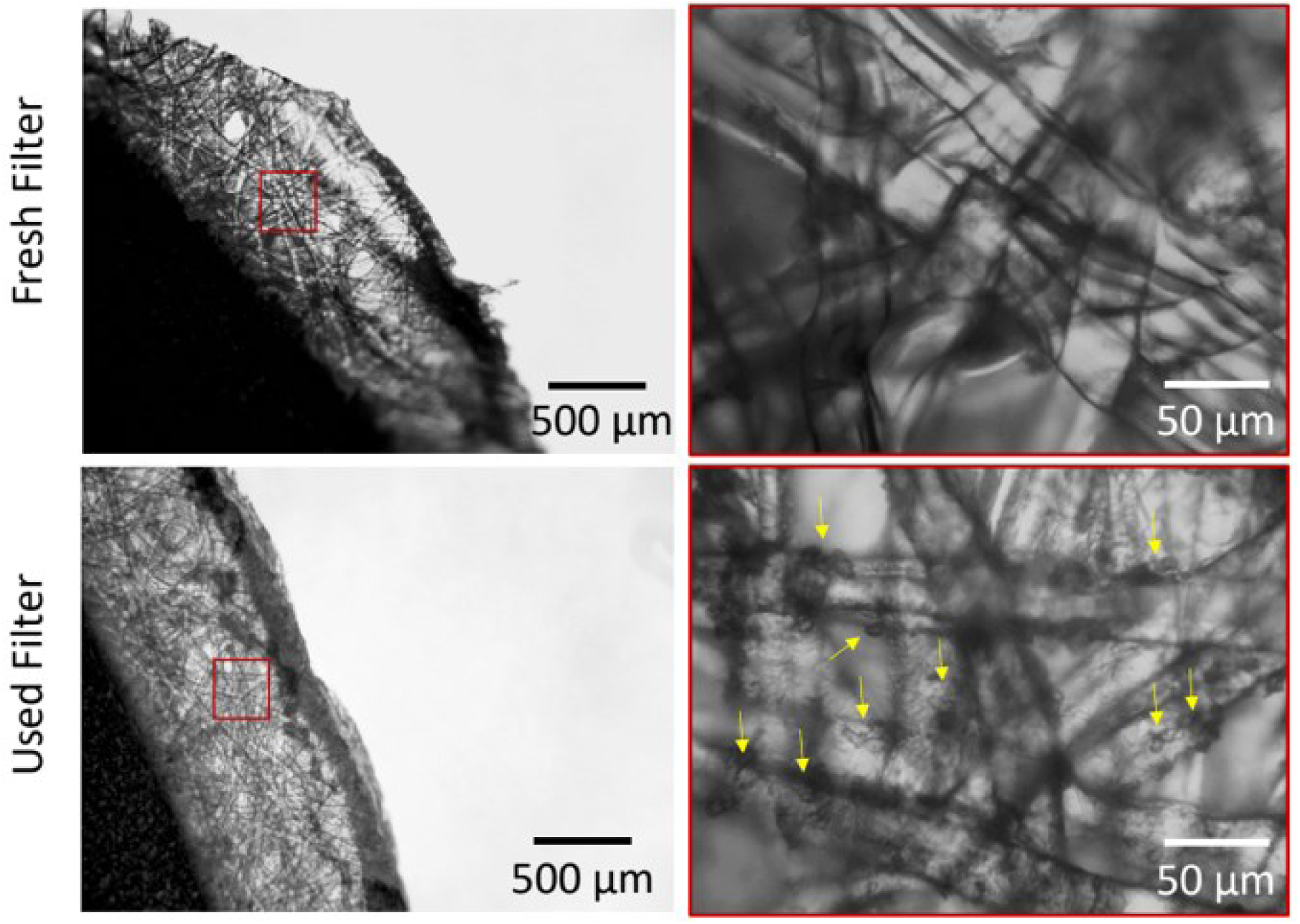
Filter membranes show visible material deposition following nanobubble filtration. Representative optical microscopy images of 5 µm PVDF filter membranes before (fresh) and after use for nanobubble filtration, acquired at multiple magnifications. Used membranes show visible accumulation of deposited material on the membrane surface and within the pore network relative to unused controls.

Furthermore, X-ray photoelectron spectroscopy (XPS) was used to examine the elemental composition of the membrane surfaces before and after use (Figure 9). XPS identifies elements at the membrane surface from characteristic core-level photoelectron peaks, labeled by element and orbital (e.g., F1s and C1s denote fluorine and carbon signal from the 1s orbital, and Na1s and P2p denote sodium and phosphorus signals from the 1s and 2p orbitals, respectively); the relative abundance of each peak reflects the relative surface concentration of that element. Comparing the two unused (fresh) filter types, the 0.45 µm membrane showed higher F1s and lower C1s abundance than the 5 µm membrane. Both membranes are manufactured from the same nominal material (hydrophilic Durapore® PVDF, per manufacturer specifications), so this baseline difference likely reflects pore-size-dependent variation in the proprietary hydrophilic surface treatment or membrane processing applied to each product line, rather than a difference in base polymer chemistry, and is unrelated to nanobubble exposure. (The manufacturer was contacted directly regarding this formulation difference but indicated that the relevant surface treatment details are proprietary and not disclosed.) Following NB filtration, however, both membrane types showed a consistent shift: Na1s and P2p abundance increased relative to their respective fresh controls, consistent with deposition of buffer salts and phospholipid headgroups on the membrane surface, while F1s abundance decreased for both filters after use. Because F1s in the fresh membranes largely reflects the underlying PVDF polymer itself, its relative decrease after filtration is consistent with a layer of retained material, phospholipid and buffer residue, coating and partially masking the native membrane surface, consistent with the optical microscopy. Notably, phosphorus was detected only on used membranes, and P2p exceeded Na1s on both membrane types, even though PBS contains roughly an order of magnitude more sodium than phosphorus. This enrichment of phosphorus relative to sodium is not what buffer salt deposition alone would produce and is consistent with phospholipid being the dominant deposit on the membrane surface, although P2p and Na1s are not exclusive to phospholipid.

**Figure 9.**
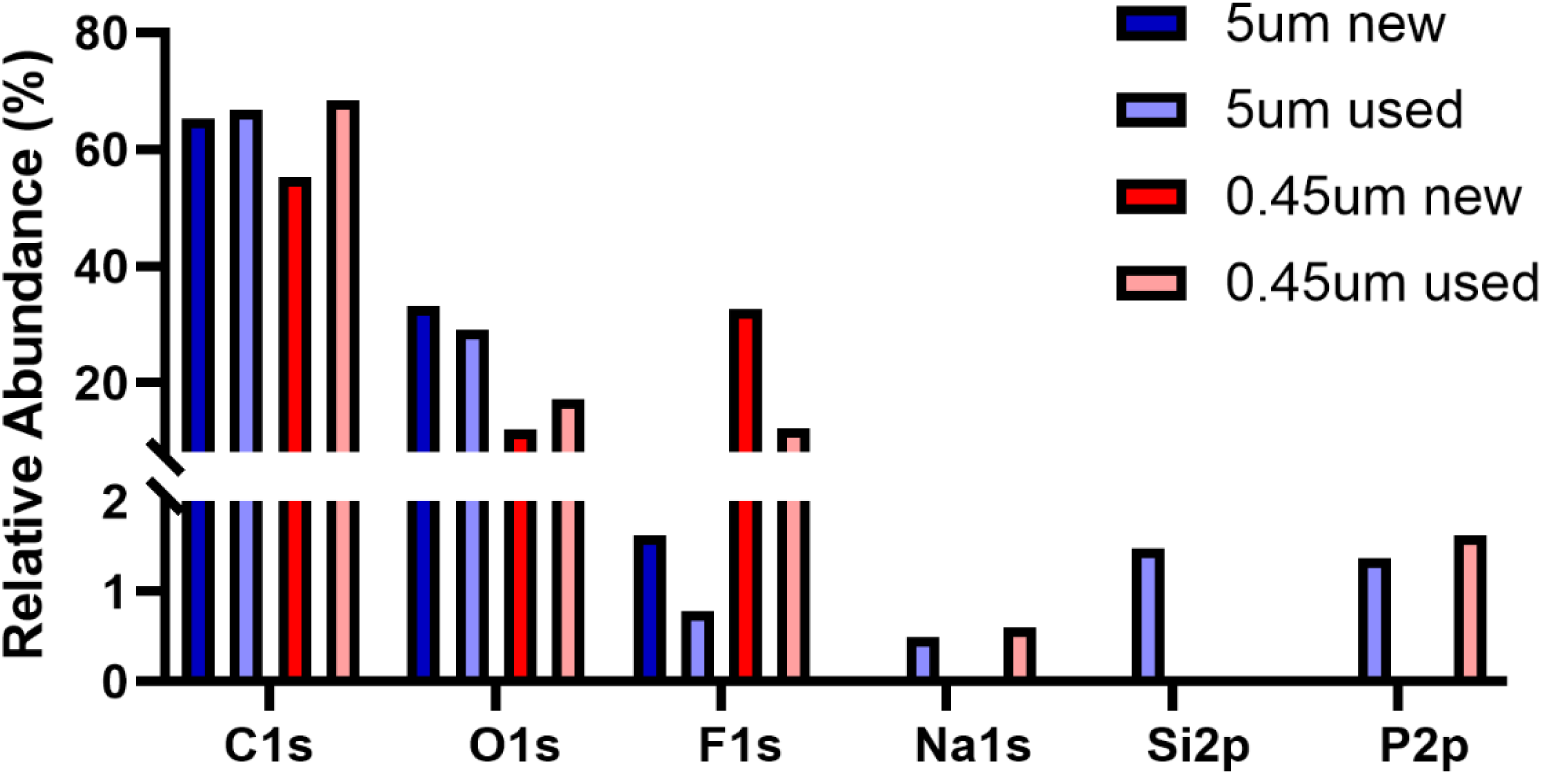
X-ray photoelectron spectroscopy reveals phospholipid deposition on filter membranes following nanobubble filtration. Relative elemental surface composition (F1s, C1s, Na1s, P2p) of 0.45 µm and 5 µm PVDF filter membranes, fresh versus after nanobubble filtration, determined by XPS. Fresh 0.45 µm membranes show higher F1s and lower C1s abundance than fresh 5 µm membranes, consistent with pore-size-dependent differences in membrane surface treatment rather than nanobubble exposure. Following filtration, both membrane types show increased Na1s and P2p abundance and decreased F1s abundance relative to their respective fresh controls, consistent with deposition of buffer salts and phospholipid headgroups on the membrane surface.

### Encapsulated gas loss tracks the loss of large bubbles common to both isolation methods, isolating shell lipid stripping as a filtration-specific effect

We next asked whether the size-dependent loss of large, gas-rich bubbles alone could account for the reduced total encapsulated gas content of filtered and centrifuged populations, or whether filtration additionally removes gas independent of size exclusion, mirroring the comparison already made for acoustic response. Encapsulated octafluoropropane (C₃F₈) content was quantified by headspace GC-MS for all four size-reduced populations relative to SOP NBs (Figure 10). Among the centrifuged populations (Figure 10B), total gas content decreased by 26% for IC-NB-100 and by 53% for IC-NB-150. Among the filtered populations (Figure 10A), gas content decreased by 18% for F-NB-5 and by 54% for F-NB-0.45.

**Figure 10.**
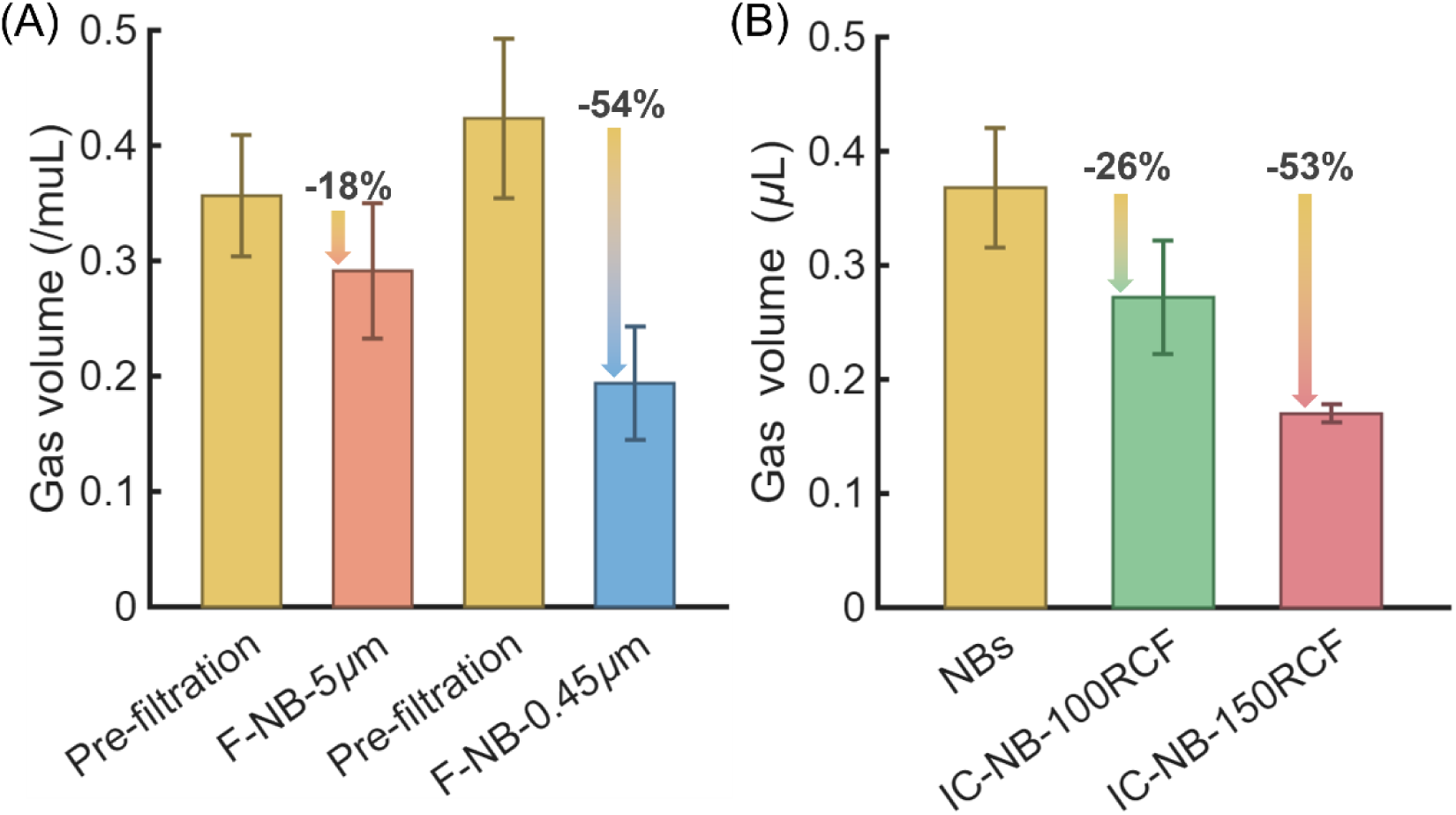
Encapsulated gas loss is comparable between size-matched filtered and centrifuged nanobubbles, dissociating gas loss from filtration-specific shell damage. Total encapsulated octafluoropropane (C₃F₈) content, quantified by headspace GC-MS and expressed as percent reduction relative to SOP NBs. **(A)** Filtered populations: total C₃F₈ volume before and after filtration for F-NB-5 (18% reduction) and F-NB-0.45 (54% reduction. **(B)** Increased-centrifugation populations: total C₃F₈ volume for SOP NBs, IC-NB-100 (26% reduction), and IC-NB-150 (53% reduction). The size-matched pair F-NB-0.45 and IC-NB-150 (both truncated near 400 nm, Figure 1) shows near-identical gas loss (54% vs. 53%), indicating that gas loss in both isolation methods is driven predominantly by removal of the largest, most gas-rich bubbles rather than by the isolation method itself. Data are shown as mean ± SD, n = 3 independent batches per group;

**Figure 11.**
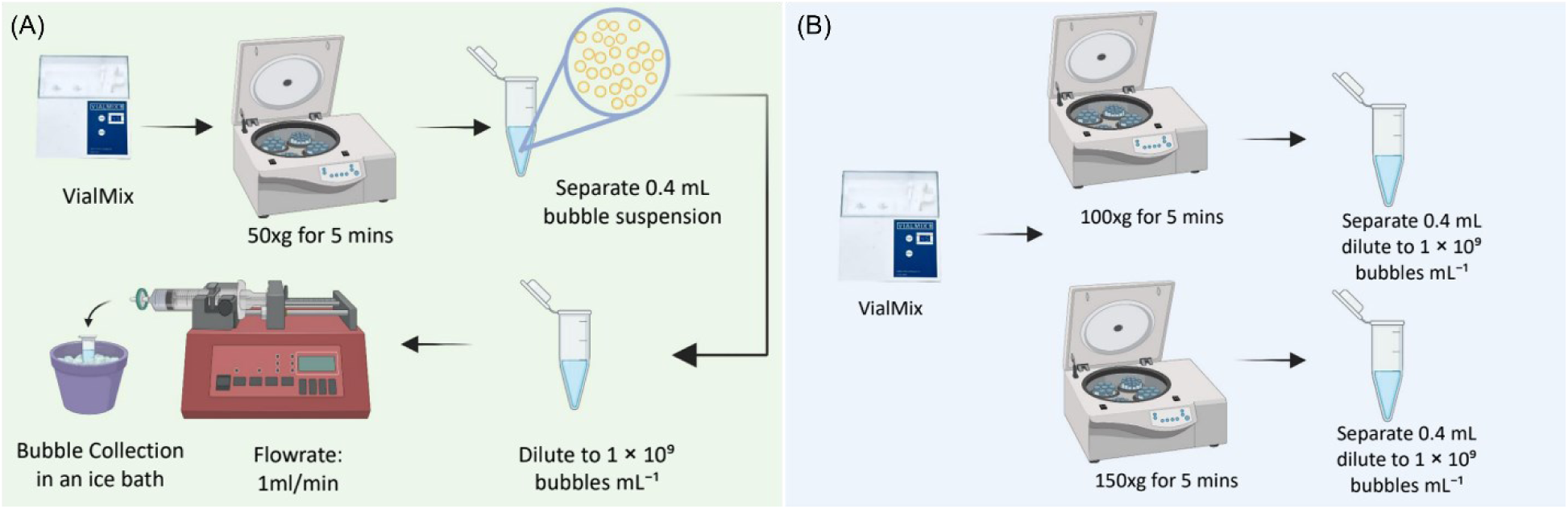
Size-isolation strategies for nanobubble (NB) suspensions. **(A)** Schematic of membrane-based size isolation, in which buoyant NB suspensions were passed through PVDF syringe filters with nominal pore sizes of 5.0 or 0.45 µm at a constant flow rate of 1.0 mL min^-1^. The 5.0-µm filter was used to assess the effect of membrane processing in the absence of substantial size exclusion, whereas the 0.45-µm filter imposed a more stringent size cutoff. **(B)** Schematic of differential centrifugation, in which freshly activated bubble suspensions were centrifuged at progressively higher relative centrifugal forces (RCF; 50, 100, and 150 × g) for 5 min. The 50 × g buoyant fraction was used as the SOP NB reference, while increased centrifugation generated progressively narrower NB populations (IC-NB-100 and IC-NB-150) without membrane contact. The two approaches enable comparison of size reduction achieved by physical size exclusion versus differential buoyant separation.

Notably, the two size-matched populations, F-NB-0.45 and IC-NB-150, both truncated near 400 nm (Figure 1), showed nearly identical gas loss (54% versus 53%, respectively). Because gas volume scales with the cube of bubble radius, removal of the upper size tail is expected to produce a disproportionate reduction in total encapsulated gas regardless of how that removal is achieved; the close agreement between F-NB-0.45 and IC-NB-150 indicates that this size-dependent effect accounts for the majority of the observed gas loss in both cases, and that filtration does not itself directly remove encapsulated gas beyond what centrifugation-based size reduction alone would predict. Any additional gas loss caused by filtration-induced shell destabilization would be expected to arise indirectly, through accelerated diffusion of gas across a compromised shell, rather than as a direct mechanical consequence of the filtration process. A similar, more modest pattern was observed for F-NB-5 and IC-NB-100 (18% and 26%, respectively), consistent with the comparatively minor size-distribution truncation associated with each of these separations.

This result is important for interpreting the lipid and shell-integrity data above. If filtration destabilized bubbles primarily by depleting encapsulated gas, filtered populations would be expected to show excess gas loss relative to size-matched centrifuged controls; instead, gas loss is essentially indistinguishable between the two methods once size distribution is matched. Using an entirely independent method, GC-MS thus gives no evidence of excess gas loss as the basis of the acoustic difference, consistent with F-NB-0.45 particles remaining gas-filled bubbles. (Figures 6 and 7): F-NB-0.45 particles are still genuine, gas-filled nanobubbles, retaining essentially the same total C₃F₈ content as their size-matched IC-NB-150 counterparts, rather than collapsed remnants or non-gaseous debris. What differs between the two populations is not whether gas is present, but the integrity and composition of the lipid shell that surrounds it: F-NB-0.45 carries a shell measurably depleted of lipid beyond what free-lipid removal or size exclusion can explain (Figures 6–7), leaving a less mechanically robust coating around an otherwise intact gas core. This shell deficit, rather than gas loss, is the most parsimonious explanation for why F-NB-0.45 is destroyed by ultrasound exposure under which its size- and gas-matched IC-NB-150 counterpart remains stable (Figures 2 to 5).

## Discussion

This study addresses a specific methodological assumption underlying much of the current debate over nanobubble echogenicity: that size-fractionation procedures broadly, and membrane filtration in particular, act as passive tools that isolate a submicron population without altering its physical properties. Across acoustic and compositional assays, filtered and centrifuged nanobubbles with matched size distributions behaved differently, and this divergence tracked with membrane exposure rather than with bubble size itself. Size did contribute to the acoustic response: IC-NB-150 produced roughly 45–55% less signal than the larger centrifuged populations, as expected from reduced scattering. However, this decrease was far smaller than the complete loss of signal in F-NB-0.45, whose signal was indistinguishable from PBS. Gas-content measurements further indicated that filtered bubbles are not simply destroyed or emptied during processing; they persist as gas-core particles, but with lower total lipid than centrifuged bubbles, which is not fully accounted for by removal of free lipid. Together, these observations are consistent with a change in shell lipid content and mechanical robustness, rather than bubble elimination, contributing to the reduced acoustic activity seen after filtration. A reduction in shell lipid content would be expected to lower the lipid coverage of the bubble surface, which in shell models such as the Marmottant model can shift a bubble from the buckled toward the ruptured regime and change its acoustic response; modeling of this effect will be presented in a separate future study.

These conclusions draw on complementary measurements of size (RMM, NTA, DLS), gas content (GC-MS), lipid content (Stewart assay, RMM), and membrane deposition (optical microscopy, XPS). The size and gas data indicate that F-NB-0.45 and IC-NB-150 are comparable gas-filled populations, while the lipid and membrane data indicate that filtration removes substantially more lipid, and are consistent with deposition of some of this material on the membrane. No single measurement here isolates shell lipid directly, so the shell-depletion interpretation rests on the combined pattern of these results rather than on any one assay.

This distinction has practical consequences for how nanobubble echogenicity has been interpreted in the literature. Sheth et al. [1] reported that filtering a lipid-shelled C₃F₈ nanobubble suspension through a 0.45 µm PVDF membrane reduced the mean bubble diameter, the fraction of bubbles larger than 1 µm, and the ultrasound signal, and concluded that the apparent echogenicity of nanobubble suspensions arises from pre-existing microbubbles that sizing methods do not reliably detect. Our results are consistent with larger bubbles contributing to the signal of the unfiltered suspension, and we do not dispute that filtration removes them. They indicate, however, that filtration also changes the remaining population, so a loss of signal after filtration cannot be attributed to the removed size fraction alone. The two studies used the same shell composition, gas, centrifugation protocol, and membrane material, so membrane-induced shell changes may also have occurred in the earlier work. How large they were would depend on the filter format, flow rate, and dilution, which are not reported or differ from ours (Sheth et al. matched filtered and unfiltered samples by total bubble volume and imaged at 6–7 MHz and, in vivo, at 18 MHz, whereas we filtered at 1.0 mL min^−1^ and imaged at 21 MHz), and we have not tested their exact conditions. Sheth et al. also note that gas content was not measured in their study and that GC-MS would help assess the effect of filtration on bubble destruction; the gas-content and lipid measurements reported here address that point for our system. Conversely, our data do not exclude a contribution from residual larger bubbles in the centrifuged populations, since NTA and DLS can miss rare large particles. Studies that use filtration to narrow a size distribution, including work that has treated post-filtration populations as a size-controlled baseline for isolating the acoustic contribution of bubble geometry, implicitly assume that any change in signal following filtration reflects the removed size fraction rather than a change in the remaining population. Our results suggest this assumption may not hold for lipid-shelled formulations, and that conclusions drawn from filtered nanobubble populations, whether supporting or challenging nanobubble echogenicity, may be confounded by processing-induced shell changes. This does not resolve the underlying controversy over whether submicron bubbles can generate clinically relevant contrast, but it does indicate that the isolation method used to reach a given size distribution is itself an experimental variable that warrants explicit reporting and control, rather than an incidental detail of sample preparation.

A simple mass-balance argument supports this interpretation. If filtration acted only by removing larger bubbles, the extra lipid lost by the 0.45 µm filter should match the lipid those bubbles carried. Using a monolayer estimate, the removed bubbles carried only ∼0.0003 mg, about 5% of the extra loss, which makes size exclusion an insufficient explanation. The residual is consistent with lipid being stripped from shells of surviving bubbles and deposited on the membrane (Figs. 8–9), though it may also include free lipid below the RMM detection limit. The shell-depletion interpretation therefore rests on elimination of the alternatives plus the acoustic data, not on a direct shell measurement.

The selective loss of shell lipid without a proportional loss of encapsulated gas is also informative about the physical process occurring at the membrane. Lipid monolayers stabilizing gas nanobubbles are under interfacial tension and can undergo local buckling, desorption, or reorganizing when subjected to shear or elongational stress, such as that experienced during passage through a constricted pore [25]. To estimate the magnitude of this stress, flow through each membrane was modeled as pressure-driven flow through an idealized bundle of cylindrical pores, using the pore diameter, membrane thickness, porosity, and flow rate used experimentally. Because real membranes are more tortuous and offer greater flow resistance than an idealized cylindrical geometry, this approach provides a conservative, lower-bound estimate of the true local stress at the pore. Even under this conservative model, the 0.45 µm membrane subjected bubbles to a wall shear stress and wall shear rate approximately 11-fold higher than the 5 µm membrane (0.86 Pa and 862 s^−1^, versus 0.078 Pa and 78 s^−1^, respectively), consistent with the substantially greater lipid depletion and instability observed for F-NB-0.45 relative to F-NB-5. Notably, the confinement ratio (bubble diameter relative to pore diameter, assuming a nominal bubble diameter of 350 nm) remained below 1 for both membranes (0.78 for the 0.45 µm filter and 0.07 for the 5 µm filter), indicating that a typical bubble is smaller than the pore diameter for both membranes. This does not hold for the largest bubbles in the SOP input (up to ∼800 nm), which exceed the 0.45 µm pore diameter, so some geometric confinement is expected for that filter. The difference in outcome between the two filters may therefore reflect both the higher local shear and a greater degree of confinement at the 0.45 µm pore. This idealized, single-bubble model also does not account for possible pore-mouth crowding at the bubble concentrations used experimentally, or for membrane wettability and lipid– surface electrostatic interactions, either of which could further modulate the local stress and adhesion experienced by bubbles at the pore entrance. It is because of effects like these that filter manufacturer guidelines specifically state that the filters are not appropriate for the filtration of colloidal suspensions and emulsions without validation. Such stress could preferentially remove loosely packed or excess shell material, with the displaced lipid subsequently adsorbing to the membrane surface, consistent with the well-documented tendency of phospholipid vesicles to adsorb onto hydrophilic supports and form deposited lipid layers [26], while leaving the bubble’s core structurally continuous enough to remain intact and gas filled, at least until it is subsequently challenged by an acoustic stimulus. The similarity in gas loss between size-matched filtered and centrifuged populations argues that this shell-stripping process occurs largely independent of bulk gas diffusion, consistent with a mechanically driven rather than diffusion-driven origin for the observed instability [27].

Several limitations should be considered when interpreting these findings. All experiments were performed using a single lipid shell composition and a single encapsulated gas (C₃F₈); whether the sensitivity to filtration observed here generalizes to other formulations or gas cores remains to be established. None of the characterization methods used here directly resolves the lipid composition or packing density of a given individual bubble shell. Consistent with this, the per-bubble lipid mass used here is a population-averaged quantity that combines shell-associated and co-purifying free lipid, and the monolayer-coverage estimate relies on a representative phospholipid packing area (∼0.65 nm² per headgroup [28]) and the nominal 6:1:2:1 shell stoichiometry rather than a directly measured single-bubble composition; they should therefore be read as order-of-magnitude bounds that constrain, but do not on their own resolve, the absolute lipid content of individual shells. Cryo-electron microscopy could help establish whether filtered and centrifuged shells differ in gross morphology, while resolving subtler packing differences would likely require complementary techniques such as fluorescence-based packing probes or atomic force microscopy nanoindentation. Filtration was limited to one membrane material (PVDF), two pore sizes (0.45 and 5 µm) from two product lines, and a single flow rate. Different membrane materials or processing conditions may produce different degrees of shell disruption, consistent with reports that filtration performance for lipid-based formulations varies considerably with membrane and process parameters even for conventional liposomes [29]. Testing additional membrane materials, matched product formats, and flow rates is an important next step. Increased centrifugation produced smaller statistical comparisons were based on n = 3 independent batches per group, which limits power and precludes formal tests of normality and variance homogeneity. Filtered and centrifuged populations were imaged in separate phantoms, each normalized to its own PBS control, rather than in a single shared acquisition. Also, size-matching between F-NB-0.45 and IC-NB-150 was established from RMM, NTA, and DLS distributions; because scattering is weighted toward the largest bubbles, small differences in the 350–450 nm tail that these methods cannot fully resolve could contribute to the acoustic differences observed. Finally, all measurements were performed in vitro under controlled buffer conditions, and the acoustic and mechanical stability of nanobubbles in physiological or flow environments may differ from what is reported here.

Translating these findings into practical changes for nanobubble research and manufacturing will require further work. Establishing whether the shell lipid depletion observed here generalizes across formulations and filter types would help determine whether membrane filtration should be reconsidered as a standard nanobubble isolation step, or whether it remains acceptable for some material and pore combinations but not others. Isolation strategies that avoid membrane contact, such as controlled centrifugation protocols or acoustic or field-flow fractionation, should be compared with filtration across a range of formulations to identify methods that narrow size distributions without compromising shell integrity. Membrane-based alternatives such as tangential flow filtration would also need to be evaluated directly, since they may involve similar membrane–bubble interactions. A stability metric that captures bubble robustness under mechanical or acoustic stress, rather than size and concentration alone, could also be explored as a quality-control measure for nanobubble formulations. More broadly, shell integrity may influence nanobubble behavior not only under ultrasound but also in vivo, including circulation, extravasation, and drug release. Therefore, it will be important to determine how filtration-induced shell changes affect biological performance before applying isolation protocols developed for acoustic characterization to clinical formulations.

## Materials and Methods

### Standard Nanobubble Fabrication

Perfluoropropane (C₃F₈) nanobubbles (SOP NBs) were prepared following an established protocol [30]. Briefly, a lipid mixture of 1,2-dibehenoyl-sn-glycero-3-phosphocholine (DBPC), 1,2-dipalmitoyl-sn-glycero-3-phosphate (DPPA), 1,2-dipalmitoyl-sn-glycero-3-phosphoethanolamine (DPPE), and 1,2-distearoyl-sn-glycero-3-phosphoethanolamine-N-[methoxy (polyethylene glycol)-2000] (DSPE-mPEG2000) was combined at a molar ratio of 6:1:2:1 and dissolved in 1.03 g propylene glycol (PG) at 80 °C until fully solubilized for a total lipid concentration of 10 mg mL^-1^. Preheated glycerol/PBS solution (9 mL, 0.1575 g mL^−1^ glycerol in phosphate-buffered saline) was added, and the resulting suspension was sonicated for 10 min at room temperature (CPX2800H, Branson Ultrasonics, Emerson, Danbury, CT, USA) to ensure complete lipid dissolution and homogenization.

For bubble generation, 1 mL aliquots of the lipid solution were dispensed into 3 mL borosilicate serum vials, sealed with rubber septa and aluminum crimp caps, and the vial headspace was exchanged with C₃F₈ gas. Bubbles were formed by mechanical agitation for 45 s using a clinical amalgamator (VialMix®, Bristol Myers Squibb, New York, NY, USA), yielding a suspension containing a broad, polydisperse distribution of gas-filled bubbles spanning the nano-and microscale.

Freshly activated bubble suspensions were inverted and centrifuged at 50 relative centrifugal force (RCF) for 5 min to isolate the buoyant nanobubble fraction. From each vial, 400 µL of the buoyant fraction was withdrawn using a 25G × 5/8″ hypodermic needle (BD PrecisionGlide™, Becton Dickinson, Franklin Lakes, NJ, USA; Cat. No. 305122) inserted approximately 5 mm into the inverted cap. The SOP NBs served as the reference population for all comparisons. Samples were kept on ice and used immediately.

### Narrowing NB Size Distribution

Two independent strategies were used to narrow the bubble size distribution: differential centrifugation, which relies on the intrinsically slower buoyant rise of smaller bubbles, and membrane filtration, which imposes a mechanical size cutoff. These strategies were compared directly to distinguish the effect of size reduction from the effect of the isolation method itself.

#### Increased centrifugation

To generate progressively narrower size distributions without membrane contact, freshly activated vials from the same formulation were centrifuged at 100 or 150 RCF for 5 min, using the same isolation procedure as the SOP NBs. IC-NB isolates were prepared independently from the SOP isolate and were not derived from it. These populations are referred to as increased-centrifugation nanobubbles (IC-NB-100 and IC-NB-150, respectively).

#### Membrane filtration

Buoyant NB suspensions were diluted to 1 × 10⁹ bubbles mL^−1^, determined by resonant mass measurement (RMM), and passed through either 0.45 µm or 5.0 µm polyvinylidene fluoride (PVDF) syringe filters (Millex™-HV, Cat. No. SLHV033R, and Millex™-SV, Cat. No. SLSV025LS, respectively; Merck Millipore, Burlington, MA, USA) using a programmable syringe pump (Pump 11 Pico Plus Elite, Harvard Apparatus, Harvard Bioscience Inc., Holliston, MA, USA) at a constant flow rate of 1.0 mL min^−1^. The 5.0 µm pore size was selected as roughly fivefold larger than the largest bubbles present in the SOP suspension, allowing filtration effects to be assessed independently of size exclusion. Filters were not pre-wetted prior to bubble suspension filtration. Filtered samples (F-NB-0.45 and F-NB-5) were collected on ice and characterized immediately. Following filtration, filters were disassembled and membranes retained for microscopy and surface chemical analysis; unused filters of the same type served as membrane controls.

### Nanobubble Characterization

Particle size distribution and concentration were characterized using three complementary techniques. Total particle size distribution and concentration were measured by nanoparticle tracking analysis (NTA; ViewSizer 3000, HORIBA Instruments Incorporated, Particle Characterization Division, Irvine, CA, USA). Buoyant, gas-filled particle size and concentration were measured by resonant mass measurement (RMM; Archimedes Particle Metrology System, Malvern Panalytical Ltd., Malvern, UK), which distinguishes positively buoyant NBs from negatively buoyant lipid vesicles, micelles, and other non-gas-filled debris by the sign of their buoyant mass. Hydrodynamic size was additionally assessed by dynamic light scattering (DLS; Zetasizer Advance, Malvern Panalytical Ltd., Malvern, UK). The RMM nano-sensor used in this study is limited to particles below 2 µm; DLS and NTA were used to extend size characterization up to 15 µm and determine the absence or presence of a residual large-particle tail beyond the RMM detection limit.

All samples were measured within 10–15 min of preparation. Each sample was diluted in PBS to a concentration within the working range of each instrument, and the dilution factor was recorded and used to back-calculate the concentration of the undiluted sample. Unless otherwise stated, reported NB concentrations are the total particle concentration measured by NTA multiplied by the buoyant (gas-filled) fraction determined by RMM, which restricts the reported values to gas-filled particles. Because RMM detects buoyant particles only above its lower size limit (∼100 nm), this concentration excludes bubbles below that size. For the filtration, ultrasound, and lipid-assay experiments, SOP NB and IC-NB samples were diluted to 1 × 10⁹ NB mL^−1^. Aliquots of the diluted SOP suspension were passed through the syringe filters to generate the F-NB samples. F-NB samples were characterized and imaged immediately after filtration, without further dilution or concentration.

### Headspace Gas Chromatography–Mass Spectrometry (GC-MS)

Encapsulated C₃F₈ content was quantified before and after filtration using headspace GC-MS. Aliquots (500 µL) of NB suspension were sealed in GC-MS headspace vials and heated in a 50 °C ultrasonic bath (CPX2800H, Branson Ultrasonics) for 20 min to release encapsulated gas into the vial headspace.

Headspace samples (5 µL) were analyzed using a 7890B gas chromatograph coupled to a 5977B mass selective detector (Agilent Technologies, Santa Clara, CA, USA), introduced via autosampler at a split ratio of 5:1 with the injector held at 200 °C. Separation was performed on an HP-5MS capillary column (30 m × 250 µm × 0.25 µm; Agilent Technologies) using helium carrier gas at 1.5 mL min^−1^. The oven program held 60 °C for 1 min, ramped at 30 °C min^−1^ to 120 °C, and held for 5 min; the transfer line was maintained at 250 °C.

Detection was performed in selected ion monitoring (SIM) mode (electron multiplier gain factor of 2), monitoring characteristic C₃F₈ fragment ions at m/z 69 (CF₃⁺) and m/z 169 (C₂F₅⁺) with a 50 ms dwell time. Fragment identity was confirmed against the NIST mass spectral database, and quantification was based on the dominant CF₃⁺ fragment (retention time ≈ 4.48 min) using a calibration curve constructed from known C₃F₈ standards to convert peak area to equivalent gas volume.

### Lipid Quantification

Total phospholipid content before and after filtration was quantified using the Stewart ferrothiocyanate assay. Pre-filtration NB suspensions were diluted to approximately 1 × 10⁹ bubbles mL^−1^, and 2.5 mL aliquots were frozen at −80 °C and lyophilized to complete dryness. Lipids were extracted from the freeze-dried material with 2.0 mL chloroform (CHCl₃). Following extraction, the clear chloroform-soluble fraction was carefully separated from the insoluble residue. The recovered chloroform fraction was adjusted to a final volume of 2.0 mL with chloroform, followed by the addition of 2.0 mL of ammonium ferrothiocyanate reagent. Samples were vortexed for 1 min (VWR International, Radnor, PA, USA) and allowed to phase-separate; the chloroform phase containing the complex was collected for spectrophotometric analysis.

Absorbance spectra (200–800 nm, 1 nm resolution) were acquired on a Spark multimode microplate reader controlled by SparkControl software v3.2 (Tecan Trading AG, Männedorf, Switzerland). Lipid concentration was calculated from the absorbance peak at 485–488 nm using calibration curves generated from lipid standards processed under identical conditions.

Because the Stewart assay cannot distinguish shell-associated lipid from free (non-bubble-associated) lipid aggregates, this measurement was complemented by RMM quantification of the negatively buoyant particle fraction (free lipid/liposomal material) before and after filtration, allowing shell-derived lipid loss to be estimated independently (see Results).

### Estimation of bubble-associated lipid from RMM size distributions

To estimate how much of the lipid lost during filtration could be attributed to the removal of larger bubbles, the shell lipid mass of a bubble of radius *r* was approximated assuming a single lipid monolayer. The shell mass was calculated as 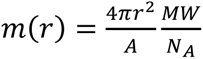, where *A* is the area occupied per lipid molecule, *MW* is the molar-mass-weighted mean molecular weight of the DBPC:DPPA:DPPE:DSPE-mPEG2000 (6:1:2:1) formulation, and *N_A_* is Avogadro’s number. Using a mean molecular weight of 1027 g mol^−1^ and an assumed molecular area of 0.50 nm² per lipid, representative of a condensed lipid monolayer, the shell mass is given by *m*(*r*) = 4.3 × 10^−5^*r*^2^fg, where *r* is the bubble radius in nm [28]. The buoyant-particle concentration per size bin (RMM, mean of n = 3) of F-NB-5 was subtracted from that of F-NB-0.45 bin by bin to give a difference distribution, representing the net bubbles present after 5 µm but absent after 0.45 µm filtration (whether retained or destroyed on the membrane). Concentrations were scaled to the 2.5 mL volume used for the Stewart assay. The lipid mass attributable to this population was obtained as Σᵢ *Nᵢ* · *m*(*rᵢ*) over all bins. This is an upper-bound estimate of bubble-associated lipid, since the RMM detects buoyant particles only above ∼100 nm, and it was compared with the difference in total lipid measured by the Stewart assay between the two samples. The calculation was repeated with *m*(*r*) scaled up to [10×/22×] to test sensitivity to the shell-mass assumption.

### Filter Membrane Characterization

To assess physical and chemical evidence of nanobubble–membrane interaction, both unused and NB-exposed filter membranes were analyzed by optical microscopy and X-ray photoelectron spectroscopy (XPS). Following filtration, membranes were removed from their housings and dried in a desiccator prior to analysis.

Membrane surface morphology was imaged by optical microscopy at multiple magnifications; used membranes were compared directly against unused controls of the same filter type to identify evidence of surface deposition or structural change.

Elemental surface composition was determined by XPS (PHI VersaProbe III Scanning X-ray Photoelectron Spectrometer, ULVAC-PHI, Chigasaki, Japan) using a monochromatic Al Kα source and a 200 × 200 µm analysis area. Survey and high-resolution spectra were acquired and analyzed using the manufacturer’s software, with binding energies referenced to the C1s peak at 284.5 eV; relative elemental composition was calculated from peak areas following standard sensitivity-factor correction.

### Ultrasound Response Characterization

Acoustic response of F-NBs and IC-NBs was evaluated using a Vevo 2100 high-frequency ultrasound imaging system with an MS250 linear array transducer (21 MHz nominal center frequency; FUJIFILM VisualSonics Inc., Toronto, ON, Canada). Samples were loaded into individual wells of a custom four-well phantom cast from 1.5% (w/v) agarose, and imaging was initiated only after all wells had been loaded to ensure simultaneous, time-matched acquisition across samples. SOP NBs and IC-NB-100/150 suspensions were each diluted to 1 × 10⁹ NB mL^−1^, determined by NTA and RMM, prior to imaging. F-NB-0.45 and F-NB-5 were generated by passing an aliquot of this same 1 × 10⁹ NB mL^−1^ SOP suspension through the 0.45 µm or 5 µm filters and were imaged directly at their resulting post-filtration concentration, without further dilution or re-concentration. Concurrent B-mode and contrast-mode images were acquired at 1 frame s^−1^ for 300 frames at each of three acoustic pressure settings.

Raw in-phase and quadrature (IQ) data were exported for offline analysis in MATLAB (MathWorks, Natick, MA, USA). Regions of interest corresponding to each well were analyzed independently to extract acoustic signal amplitude, which was normalized to a PBS control acquired simultaneously under identical imaging conditions in the same phantom. Temporal signal evolution across the 300-frame acquisition was used to assess acoustic persistence and susceptibility to ultrasound-induced bubble destruction.

### Statistical Analysis

Experiments were performed using three independently prepared NB batches per condition (n = 3), where each batch was prepared separately beginning at mechanical amalgamation from a stock lipid dispersion. The independent batch was the unit of replication for all statistical tests. Data are reported as mean ± standard deviation (SD) unless otherwise noted.

For ultrasound experiments, the signal amplitude of each well was normalized to the PBS control imaged simultaneously in the same phantom. For each batch, normalized amplitude was averaged across all 300 frames to give one time-averaged value per batch (B-mode and contrast mode analyzed separately). Individual frames were not treated as independent observations, because consecutive frames from the same well are serially correlated and treating them as replicates would artificially inflate the degrees of freedom and produce spuriously small p-values. Time-averaging therefore reduced each sample to n = 3 independent values.

Group differences in mean normalized amplitude were assessed by one-way analysis of variance (ANOVA), with sample type as the single factor, followed by Tukey’s honestly significant difference (HSD) post-hoc test for all pairwise comparisons, which controls the family-wise error rate across comparisons. The F-NB groups (F-NB-0.45, F-NB-5) and IC-NB groups (IC-NB-100, IC-NB-150) were compared with SOP NBs and PBS within a single six-group model so that comparisons between filtered and centrifuged populations (e.g., F-NB- 0.45 vs IC-NB-150) were corrected within the same family. Because filtered and centrifuged populations were imaged in separate phantoms on separate acquisitions, each normalized to its own simultaneously imaged PBS control, cross-experiment comparisons rely on this PBS normalization. Normalized values are dimensionless, with PBS defined as 1.

## Acknowledgments

This work was supported by the National Institutes of Health (Grant No. 5R01EB025741-07). The authors thank the members of the Kolios Laboratory (Toronto Metropolitan University) and the Exner Laboratory (Case Western Reserve University) for their valuable discussions.

## References

[1] M. Sheth et al., “Size matters: Micro-versus nanobubbles in ultrasound imaging and therapy,” Sci. Adv., vol. 11, no. 29, pp. eads2177-, 2025, doi: 10.1126/sciadv.ads2177.

2. “ISO 20480-1:2017(en), Fine bubble technology — General principles for usage and measurement of fine bubbles — Part 1: Terminology.” Accessed: Sep. 24, 2026. [Online]. Available: https://www.iso.org/obp/ui/#iso:std:iso:20480:-1:ed-1:v1:en

[3] L. E. Chen, P. Nittayacharn, and A. A. Exner, “Progress and potential of nanobubbles for ultrasound-mediated drug delivery,” Expert Opin. Drug Deliv., vol. 22, no. 7, pp. 1007– 1030, 2025, doi: 10.1080/17425247.2025.2505044.

[4] D. Wegierak et al., “Nanobubble Contrast Enhanced Ultrasound Imaging: A Review,” Wiley Interdiscip. Rev. Nanomed. Nanobiotechnol., vol. 16, no. 6, pp. e2007-n/a, 2024, doi: 10.1002/wnan.2007.

[5] A. A. Exner and M. C. Kolios, “Bursting microbubbles: How nanobubble contrast agents can enable the future of medical ultrasound molecular imaging and image-guided therapy,” Curr. Opin. Colloid Interface Sci., vol. 54, p. 101463, Aug. 2021, doi: 10.1016/j.cocis.2021.101463.

[6] F. M. Berg, M. B. Cooley, T. Kosmides, L. E. Chen, R. H. Baroni, and A. A. Exner, “New Frontiers in Contrast-Enhanced Ultrasound for Cancer Imaging,” ACS Nano, vol. 20, no. 20, pp. 14327–14350, 2026, doi: 10.1021/acsnano.5c17992.

[7] D. Sharma, T. N. Petchiny, and G. J. Czarnota, “A Promising Therapeutic Strategy of Combining Acoustically Stimulated Nanobubbles and Existing Cancer Treatments,” Cancers, vol. 16, no. 18, pp. 3181-, 2024, doi: 10.3390/cancers16183181.

[8] J. Z. Myers, J. A. Navarro-Becerra, and M. A. Borden, “Nanobubbles are Non-Echogenic for Fundamental-Mode Contrast-Enhanced Ultrasound Imaging,” Bioconjug. Chem., vol. 33, no. 6, pp. 1106–1113, 2022, doi: 10.1021/acs.bioconjchem.2c00155.

[9] N. de Jong, M. Emmer, A. van Wamel, and M. Versluis, “Ultrasonic characterization of ultrasound contrast agents,” Med. Biol. Eng. Comput., vol. 47, no. 8, pp. 861–873, 2009, doi: 10.1007/s11517-009-0497-1.

[10] M. Versluis, E. Stride, G. Lajoinie, B. Dollet, and T. Segers, “Ultrasound Contrast Agent Modeling: A Review,” Ultrasound Med. Biol., vol. 46, no. 9, pp. 2117–2144, 2020, doi: 10.1016/j.ultrasmedbio.2020.04.014.

[11] C. Chen et al., “The unique second wave phenomenon in contrast enhanced ultrasound imaging with nanobubbles,” Sci. Rep., vol. 12, no. 1, pp. 13619–12, 2022, doi: 10.1038/s41598-022-17756-1.

[12] M. B. Cooley et al., “Real-time imaging of nanobubble ultrasound contrast agent flow, extravasation, and diffusion through an extracellular matrix using a microfluidic model,” Lab. Chip, vol. 23, no. 15, pp. 3453–3466, 2023, doi: 10.1039/d3lc00514c.

[13] A. de Leon et al., “Contrast Enhanced Ultrasound Imaging by Nature-Inspired Ultrastable Echogenic Nanobubbles,” Nanoscale, vol. 11, no. 33, pp. 15647–15658, Sep. 2019, doi: 10.1039/c9nr04828f.

[14] K. Ferrara, R. Pollard, and M. Borden, “Ultrasound Microbubble Contrast Agents: Fundamentals and Application to Gene and Drug Delivery,” Annu. Rev. Biomed. Eng., vol. 9, no. 1, pp. 415–447, 2007, doi: 10.1146/annurev.bioeng.8.061505.095852.

[15] J. A. Feshitan, C. C. Chen, J. J. Kwan, and M. A. Borden, “Microbubble size isolation by differential centrifugation,” J. Colloid Interface Sci., vol. 329, no. 2, pp. 316–324, Jan. 2009, doi: 10.1016/j.jcis.2008.09.066.

[16] T. Segers, E. Gaud, M. Versluis, and P. Frinking, “High-precision acoustic measurements of the nonlinear dilatational elasticity of phospholipid coated monodisperse microbubbles,” Soft Matter, vol. 14, no. 47, pp. 9550–9561, Dec. 2018, doi: 10.1039/C8SM00918J.

[17] B. Helfield, B. Leung, X. Huo, and D. Goertz, “Scaling of the viscoelastic shell properties of phospholipid encapsulated microbubbles with ultrasound frequency,” Ultrasonics, vol. 54, Apr. 2014, doi: 10.1016/j.ultras.2014.03.014.

[18] C. C. Church, “The effects of an elastic solid surface layer on the radial pulsations of gas bubbles,” J. Acoust. Soc. Am., vol. 97, no. 3, pp. 1510–1521, Mar. 1995, doi: 10.1121/1.412091.

[19] A. J. Sojahrood, H. Haghi, T. M. Porter, R. Karshafian, and M. C. Kolios, “Experimental and numerical evidence of intensified non-linearity at the micro and nano scale: The lipid coated acoustic bubble,” Phys. Fluids, vol. 33, no. 7, p. 072006, Jul. 2021, doi: 10.1063/5.0051463.

[20] A. J. Sojahrood et al., “Influence of the liquid ionic strength on the resonance frequency and shell parameters of lipid-coated microbubbles,” J. Colloid Interface Sci., vol. 664, pp. 533–538, Jun. 2024, doi: 10.1016/j.jcis.2024.01.185.

[21] M. Yaali, A. A. Exner, and M. C. Kolios, “Acoustic characterization of shell and size-engineered submicron contrast agents using attenuation measurements,” Ultrason. Sonochem., vol. 130, pp. 107884-, 2026, doi: 10.1016/j.ultsonch.2026.107884.

[22] H. Haghi, M. Yaali, A. A. Exner, and M. C. Kolios, “Experimental investigation on the effect of concentration on the resonance frequency of lipid coated ultrasonically excited microbubbles,” Ultrason. Sonochem., vol. 112, pp. 107170-, 2025, doi: 10.1016/j.ultsonch.2024.107170.

[23] A. Jafari Sojahrood et al., “Toward Precisely Controllable Acoustic Response of Shell-Stabilized Nanobubbles: High Yield and Narrow Dispersity,” ACS Nano, vol. 15, no. 3, pp. 4901–4915, Mar. 2021, doi: 10.1021/acsnano.0c09701.

[24] C. Hernandez et al., “Sink or float? Characterization of shell-stabilized bulk nanobubbles using a resonant mass measurement technique †Electronic supplementary information (ESI) available: Experimental details, supporting information. See DOI: 10.1039/c8nr08763f,” Nanoscale, vol. 11, no. 3, pp. 851–855, Jan. 2019, doi: 10.1039/c8nr08763f.

[25] P. Marmottant et al., “A model for large amplitude oscillations of coated bubbles accounting for buckling and rupture,” J. Acoust. Soc. Am. -J ACOUST SOC AMER, vol. 118, pp. 3499–3505, Dec. 2005, doi: 10.1121/1.2109427.

[26] R. P. Richter, R. Bérat, and A. R. Brisson, “Formation of Solid-Supported Lipid Bilayers: An Integrated View,” Langmuir, vol. 22, no. 8, pp. 3497–3505, 2006, doi: 10.1021/la052687c.

[27] B. van Elburg, A. Lassus, S. Cherkaoui, G. Lajoinie, M. Versluis, and T. Segers, “Controlling the stability of monodisperse phospholipid-coated microbubbles by tuning their buckling pressure,” J. Colloid Interface Sci., vol. 685, pp. 449–457, 2025, doi: 10.1016/j.jcis.2025.01.114.

[28] J. F. Nagle and S. Tristram-Nagle, “Structure of lipid bilayers,” Biochim. Biophys. Acta BBA -Rev. Biomembr., vol. 1469, no. 3, pp. 159–195, 2000, doi: 10.1016/S0304-4157(00)00016-2.

[29] K. Zourna, A. Iwaniec, S. Turner, N. B. Jackson, and J. H. Welsh, “Optimizing the Filtration of Liposomes Using Sterilizing-Grade Filters,” PDA J. Pharm. Sci. Technol., vol. 75, no. 2, pp. 128–140, 2021, doi: 10.5731/pdajpst.2020.011866.

[30] M. B. Cooley et al., “Formulation and activation of lipid-shelled nanobubble ultrasound contrast agents,” Nat. Protoc., Jul. 2026, doi: 10.1038/s41596-026-01411-4.

