## Supplemental Figure S1 for "Membrane Filtration Removes Lipids and Compromises the Acoustic Stability of Nanobubbles"

### Supplementary material

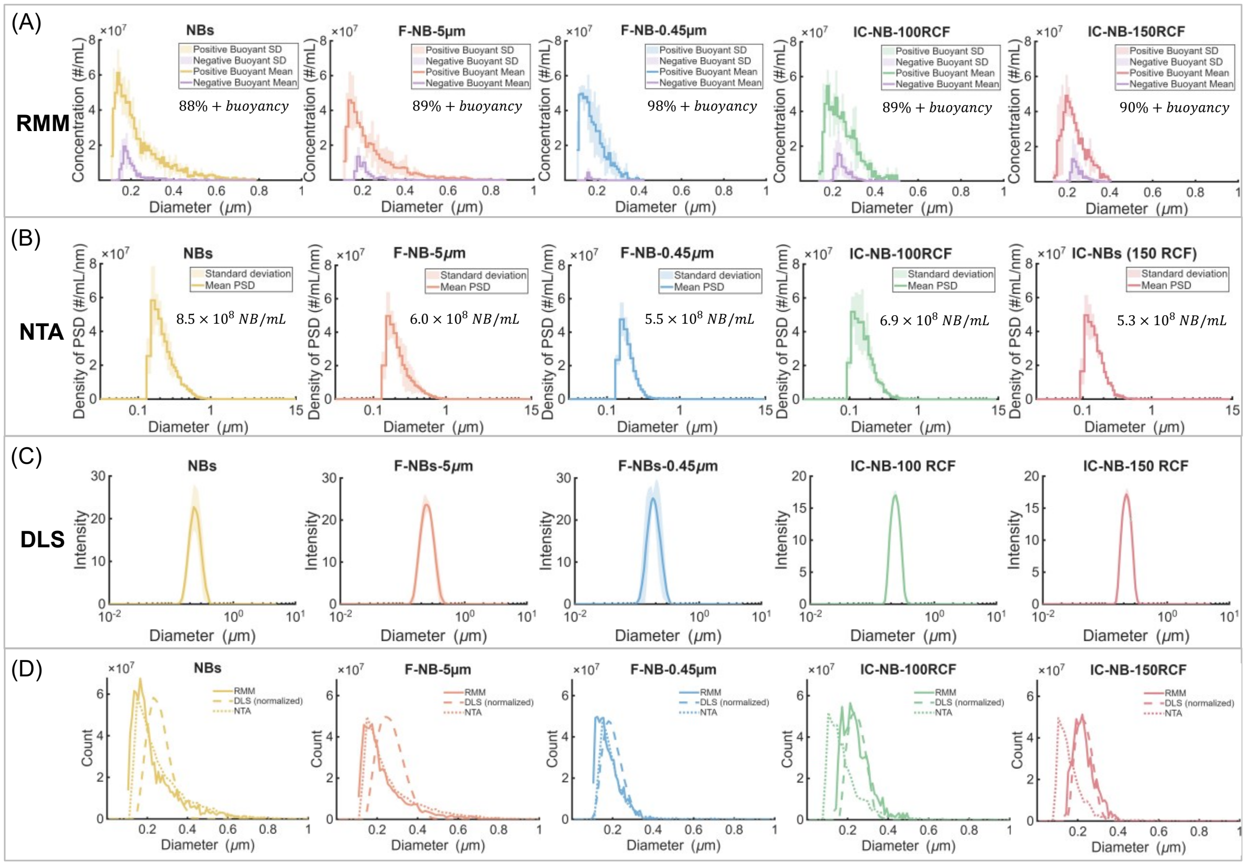


***Figure S1. Full size distribution and concentration characterization of all nanobubble populations across three measurement modalities.*** *Size distributions and concentrations for all five populations, SOP NBs (50 RCF), F-NB-5, F-NB-0.45, IC-NB-100, and IC-NB-150, measured by resonant mass measurement (RMM, top row), nanoparticle tracking analysis (NTA, second row), and dynamic light scattering (DLS, third row), with each column corresponding to one sample. Each curve represents the mean of n = 3 independent trials; shaded regions indicate SD. The RMM row is limited to the instrument's 2 μm upper detection limit, consistent with Figure 1. To resolve both the submicron peak and the absence of signal at larger sizes within a single panel, the NTA and DLS rows are plotted on a logarithmic x-axis extended to their respective upper detection limits of 15 μm and 10 μm. As summarized in Figure 1, SOP NBs and F-NB-5 show similar size distributions with an upper tail extending to approximately 800 nm, while F-NB-0.45 and IC-NB-150 are closely matched and truncated near 400 nm; IC-NB-100 shows an intermediate distribution extending to approximately 500 nm. Relative to SOP NBs, bubble concentration is reduced by 30% for F-NB-5, 35% for F-NB-0.45, 18% for IC-NB-100, and 37% for IC-NB-150. No particle population is detected above approximately 1 μm for any sample in either the NTA or DLS data, confirming that none of the isolation methods leaves a residual microbubble population capable of contributing disproportionately to acoustic scattering. The bottom row shows the mean size distributions from all three modalities overlaid for each sample, restricted to a 1 μm, linear x-axis (matching Figure 1D) for direct comparison of curve shape across modalities.*
